# Computational modeling of cancer cell metabolism along the catabolic-anabolic axes

**DOI:** 10.1101/2024.08.15.608169

**Authors:** Javier Villela-Castrejon, Herbert Levine, José N. Onuchic, Jason T. George, Dongya Jia

## Abstract

Abnormal metabolism is a hallmark of cancer. Initially recognized through the observation of aerobic glycolysis in cancer nearly a century ago. Also, we now know that mitochondrial respiration is also used by cancer for progression and metastasis. However, it remains largely unclear the mechanisms by which cancer cells mix and match different metabolic modalities (oxidative/reductive) and leverage various metabolic ingredients (glucose, fatty acids, glutamine) to meet their bioenergetic and biosynthetic needs. Here, we formulate a phenotypic model for cancer metabolism by coupling master gene regulators (AMPK, HIF-1, Myc) with key metabolic substrates (glucose, fatty acid, and glutamine). The model predicts that cancer cells can acquire four metabolic phenotypes: a catabolic phenotype characterized by vigorous oxidative processes - O, an anabolic phenotype characterized by pronounced reductive activities - W, and two complementary hybrid metabolic states - one exhibiting both high catabolic and high anabolic activity - W/O, and the other relying mainly on glutamine oxidation - Q. Using this framework, we quantified gene and metabolic pathway activity respectively by developing scoring metrics based on gene expression. We validated the model-predicted gene-metabolic pathway association and the characterization of the four metabolic phenotypes by analyzing RNA-seq data of tumor samples from TCGA. Strikingly, carcinoma samples exhibiting hybrid metabolic phenotypes are often associated with the worst survival outcomes relative to other metabolic phenotypes. Our mathematical model and scoring metrics serve as a platform to quantify cancer metabolism and study how cancer cells adapt their metabolism upon perturbations, which ultimately could facilitate an effective treatment targeting cancer metabolic plasticity.

**Statement of significance:** We present a theoretical framework that integrates both catabolic and anabolic modes of cancer metabolism, considering the complex interplay between glucose, fatty acids, and glutamine, by coupling genetic regulation with metabolic pathways. Our work characterizes four main metabolic phenotypes in cancer - OXPHOS, glycolysis, hybrid, glutamine-dominant and demonstrates the critical role of Myc on glutamine metabolism in all four phenotypes. The characterization of metabolism can guide our evaluation of patient survival across cancer types.

## Introduction

Whatever traits a cancer cell might exhibit, corresponding metabolic activities are required to support required biomass production and bioenergetic needs. Understanding cancer metabolism thus provides critical insights into various hallmarks of cancer (1), such as metastasis and immune suppression. Aerobic glycolysis, termed the Warburg effect, has been often observed in cancer. The potential benefits of aerobic glycolysis include rapid ATP synthesis, microenvironment acidification, chromatin remodeling, and increased resources available for biosynthesis (2). Initially, it was believed that this mode was necessary due to dysfunctional mitochondria. However, the last two decades have witnessed increasing evidence that mitochondrial oxidative phosphorylation (OXPHOS) plays an important role in tumorigenesis, metastasis, and drug-resistance (3). For example, murine breast cancer 4T1 cells, when entering the blood circulation, exhibit higher OXPHOS relative to the primary tumor (4). Also, Braf^*V600E*^-driven tumors exhibit increased resistance to BRAF inhibitors, accomplished in part by elevated OXPHOS and mitochondrial biogenesis (5). In short, cancer cells have been shown to actively use both glycolysis and oxidation in a manner that is dependent on their environment.

In addition to glucose, fatty acids have emerged as another important metabolic ingredient for tumorigenesis and cancer progression (6). Fatty acids support rapid tumor cell proliferation by sustaining membrane biosynthesis and in addition can serve as an important energy source during periods of metabolic stress, such as hypoxia and lipid depletion. Indeed, fatty acid oxidation (FAO) has been shown to be essential for triple negative breast cancer (TNBC) progression, the most aggressive subtype of breast cancer, regulated by oncoproteins such as SRC and Myc (7,8). Inhibiting FAO has been proposed to be a therapy for TNBC (8,9).

Another critical component of cancer metabolism is the increased consumption of glutamine (10). Glutamine fuels tumor cells through driving the tricarboxylic acid (TCA) cycle via oxidation, synthesizing fatty acids via reductive carboxylation, and giving rise to glutathione (GSH) to maintain the redox balance. One master regulator of glutamine metabolism is Myc, which can upregulate glutamine transporters to support glutamine consumption, upregulate glutaminase at both transcriptional and translational levels to promote glutamine oxidation, and drive *de novo* proline synthesis to support biosynthetic processes (11). Recent work suggests that limiting cancer progression by reducing glutamine availability can improve therapeutic outcome; however more work is required to more completely understand cancer resistance to glutamine metabolism inhibitor therapy, as well as to account for cancer heterogeneity and metabolic adaptation (12,13).

Altogether, cancer cells exhibit metabolic plasticity, which enables them to combine various metabolic ingredients and different metabolic modes to meet their biomass and energetic needs. To rationalize cancer metabolic plasticity, we initially created a model with purely genetic regulation, focusing on the interplay of two master gene regulators of OXPHOS and glycolysis - AMPK and HIF-1 (14). We showed that while normal cells in general can acquire two stable metabolic phenotypes - OXPHOS and glycolysis (when oxygen is limited), cancer cells can acquire an additional hybrid metabolic phenotype characterized by high AMPK/high HIF-1 activities, referred to as the ‘W/O’ state. We subsequently provided a more direct analysis of the ‘W/O’ state by coupling genetic regulation with three main catabolic pathways: glycolysis, glucose oxidation, and FAO (15). We showed that the ‘W/O’ state exhibits high TCA/FAO/glycolysis activity. We further showed that TNBC cells, e.g., MB-MDA-231 and SUM159, exhibit the hybrid ‘W/O’ phenotype at the population level. Therefore, dual inhibition of both OXPHOS and glycolysis in TNBC achieved the best treatment outcome (most pronounced decrease in cell proliferation and colony formation) relative to inhibiting only OXPHOS or inhibiting only glycolysis (15). This model further suggested the existence of a metabolic low/low state, characterized by low AMPK/low HIF-1 and low TCA/FAO/glycolysis. We showed that drug-tolerant melanoma cells acquired such a metabolic low/low state (16). In summary, our previous studies reveal cancer metabolic plasticity by identifying distinct phenotypes (W, O, W/O, low/low) and we have linked these phenotypes to specific gene regulators and their associated catabolic pathway activities.

In these previous efforts, we focused on the catabolic behavior of cancer cells by considering ATP-producing processes. It is clear, however, that to obtain a comprehensive understanding of cancer metabolism, we need to incorporate anabolic processes and study their interplay with the catabolic processes. In addition, as the experimental evidence of glutamine metabolism in cancer accumulates, we believe it is important to expand our approach to include various glutamine pathways. Therefore, we present here a novel metabolic network model (**Figure 1**) that explicitly considers both the catabolic and anabolic modes and includes glutamine metabolism (15). With this comprehensive metabolism model, we provide a more granular characterization of different cancer metabolic phenotypes by quantifying both catabolic and anabolic activity. Our model recapitulates the critical role of Myc in glutamine metabolism and reveals the role of glutamine metabolism in the four metabolic states. Finally, we demonstrate the functional consequence of the different metabolic states in patient survival prognosis across cancer types.

**Fig. 1:**
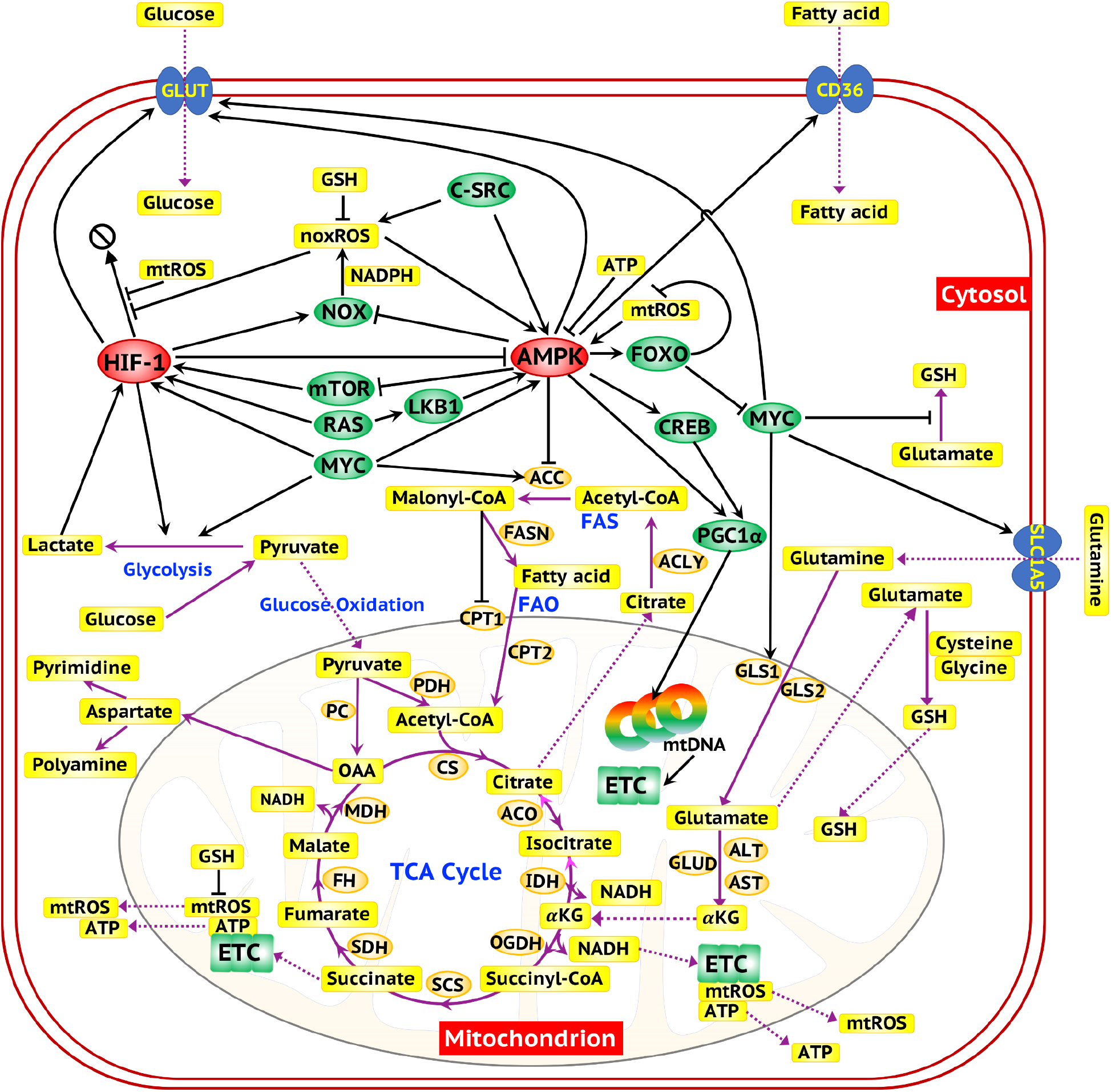
A comprehensive network featuring genetic regulation and glucose, fatty acid and glutamine metabolism in cancer. The ovals represent genes. The red ovals highlight the two master regulators of metabolism - AMPK and HIF-1. The green ovals represent oncogenes and the downstream target genes of AMPK and HIF-1. The orange ovals represent enzyme genes. The yellow rectangles represent metabolites. The black arrows represent excitatory regulatory links, and the black bar-headed arrows represent inhibitory regulatory links. The purple solid lines represent the chemical reactions involved in the metabolic pathways. The purple dotted lines represent the transportation of metabolites.

## Results

### The regulatory network of cancer metabolism

We first performed an extensive literature search (10,17,18), from which we constructed a comprehensive metabolic network featuring the uptake, transportation, and utilization of three main metabolic ingredients: glucose, fatty acid, and glutamine (**Fig. 1**). This metabolic network contains five types of regulatory interactions. *First*, metabolic pathways compete for common metabolic resources. Intracellular glucose can be used in catabolic processes in the form of glycolysis or glucose oxidation for ATP production, or alternatively in anabolic process for biomass synthesis (creating lipids, triglycerides, etc.). Intracellular fatty acids can be used in the form of FAO for ATP production, and/or anabolic processes for synthesizing the plasma membrane. Intracellular glutamine can be used in the form of glutamine oxidation for ATP production, reductive carboxylation for fatty acids synthesis, or GSH synthesis for redox balance. *Second*, the relative activities of these metabolic pathways are directly modulated by gene regulators (depicted as red or green ovals in **Figs. 1-2**). For example, HIF-1 modulates glycolysis by transcriptionally regulating expression of glycolytic enzymes. AMPK, as a key energy sensor, promotes FAO by inhibiting the lipogenic enzyme acetyl-CoA carboxylase (ACC). The oncoprotein Myc promotes glutamine oxidation via inducing glutaminase (GLS) which converts glutamine to glutamate to fuel the TCA cycle. *Third*, metabolic intermediates (depicted as rectangles in **Figs. 1-2**) can in turn affect the activities of the gene regulators. One prominent example is the reactive oxygen species (ROS). ROS, including both mitochondrial ROS (mtROS) and NADPH oxidase-derived ROS (noxROS), can stabilize HIF-1 and activate AMPK. *Fourth*, the gene regulators interact with each other. For example, AMPK and HIF-1 mutually inhibit each other. Myc post-transcriptionally enhances HIF-1. AMPK antagonizes the function of Myc through phosphorylating the transcription factor FOXO (19). *Fifth*, the gene regulators mediate the uptake of the metabolic ingredients by directly regulating the corresponding transporters. For example, HIF-1, AMPK and Myc can increase the uptake of glucose via upregulation of the glucose transporter GLUT. Myc also promotes the uptake of glutamine via upregulating the glutamine transporter SLC1A5.

### A phenotypic model of cancer metabolism

To develop a tractable mathematical model to simulate the dynamics of cancer metabolism, we coarse-grained the comprehensive metabolic network (**Fig. 1**) into a minimal network model that captures the essential features (**Fig. 2**). The minimal network model includes three main gene regulators (AMPK, HIF-1, and Myc), which modulate the uptake and utilization of the three key metabolic ingredients (glucose, fatty acid, glutamine). The minimal network model considers the dynamics of four metabolites: ROS (both mtROS and noxROS) which mediate the interplay between AMPK and HIF-1, ATP which regulates AMPK activity, acetyl-CoA which controls the inputs to the TCA cycle and GSH which modulates the cellular ROS level. The following metabolic pathways are analyzed: glycolysis, glucose oxidation, glutamine oxidation, FAO, GSH synthesis, and anabolic processes, which represent the ATP-consuming biomass-generating processes using glucose, fatty acid, and glutamine. As we will show in the following section, the minimal network model is able to capture important experimental observations about cancer metabolic plasticity.

**Fig. 2:**
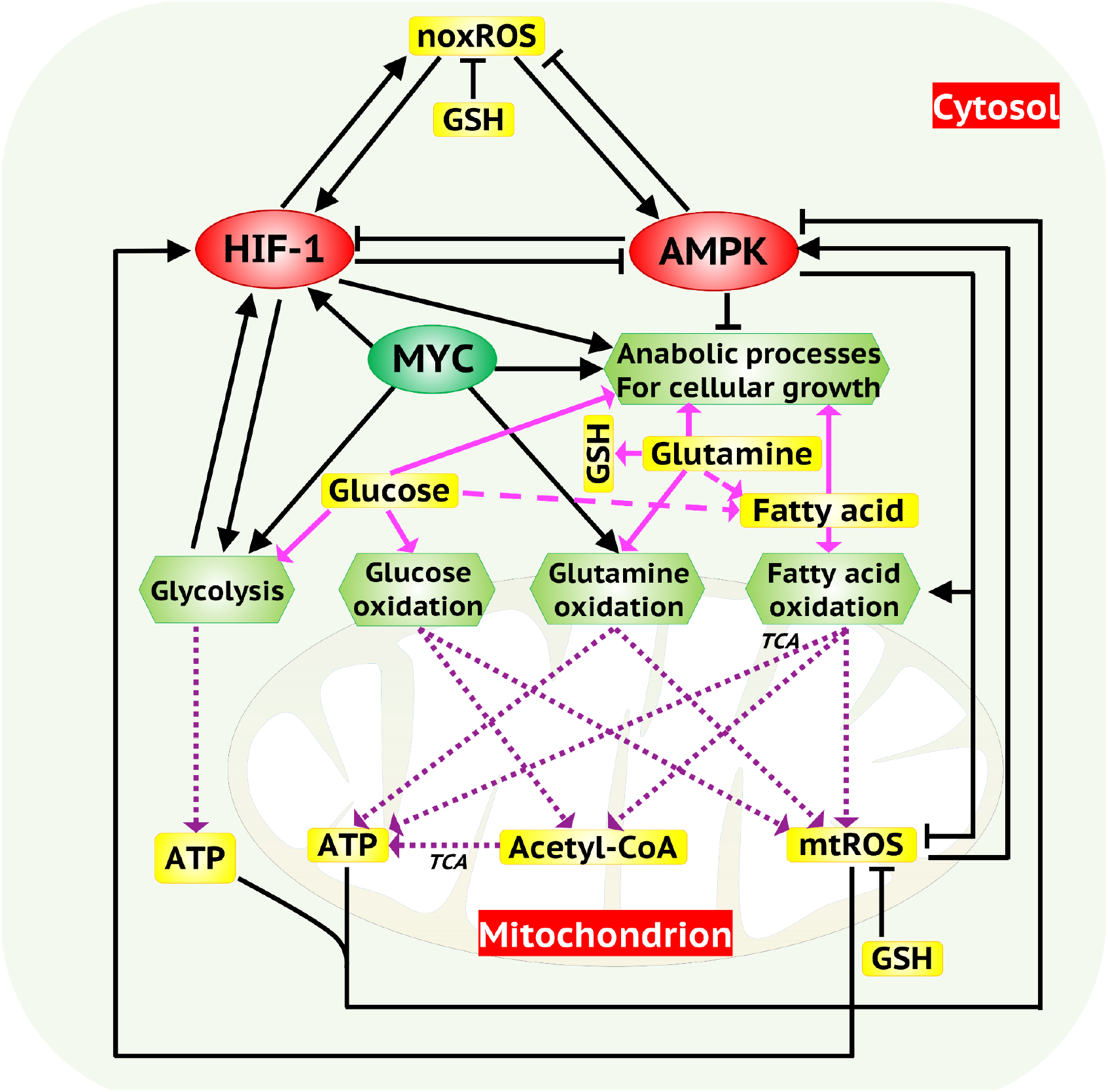
A minimal network model of cancer metabolism. AMPK, HIF-1 and Myc serve as the master regulators of cancer metabolism and regulate both catabolic processes (glycolysis, glucose oxidation, glutamine oxidation and fatty acid oxidation) and anabolic processes. The intracellular glucose can be used for glycolysis, glucose oxidation and anabolic processes. The intracellular fatty acids can be used for oxidation and anabolic processes. The intracellular glutamine can be metabolized via glutamine oxidation, synthesis of glutathione (GSH) or reductive carboxylation. The metabolites generated in the metabolic pathways, mtROS, noxROS acetyl-CoA and ATP, can in turn regulate AMPK and HIF-1. The black arrows/bar-headed arrows represent excitatory/inhibitory regulatory links. The purple dotted lines represent metabolic pathways. The magenta arrows represent the distribution of glucose, fatty acid and glutamine into different metabolic pathways. The magenta dashed lines represent fatty acid biosynthesis from glucose or glutamine.

### Cancer cells can mix and match catabolic and anabolic processes and can acquire four different metabolic phenotypes

We first identify all the possible metabolic phenotypes enabled by the regulatory network (**Fig. 2**). As overexpression of Myc has often been associated with tumor formation, we investigated how different levels of Myc affect the dynamics of the regulatory network, especially in the acquisition of different stable states (20). We discovered that when the Myc level is relatively low, cells mainly acquire two metabolic states - the Warburg state (‘W’, high HIF-1/low pAMPK) and an OXPHOS state (‘O’, high pAMPK/low HIF-1), representing the typical metabolism of normal cells (**Fig. 3A**). As the level of Myc increases, the bistability converts to tristability, and a third metabolic state - ‘W/O’ - characterized by intermediate pAMPK/HIF-1 activity emerges (**Fig. S1**). We also found that under certain conditions this network model can acquire tetra-stability (**Fig. 3B**). That is, in addition to the W, O and W/O states, the metabolic network enables cancer cells to acquire an additional stable state - the ‘Q’ state, which relies mainly on glutamine oxidation (as shown in the next section) (**Fig. 3B)**. The ‘Q” state exhibits both low pAMPK and low HIF-1 activity, and low glycolysis, glucose/fatty acid oxidation, and is analogous to our previously identified “low/low phenotype” using a simplified regulatory network without glutamine metabolism (16). We then reveal the metabolic pathway activities of the four states - W, O, W/O and Q (**Fig. 3C**). The “W” state is distinguished by heightened glycolysis activity (G2), and it exhibits increased anabolic activities, which are attributed to the reductive metabolism of glucose (Gre), fatty acids (Fre), and glutamine (Qre). The “O” state is characterized by high catabolic activities including oxidation of glucose (G1), fatty acid (F), and glutamine (Q1), and notably high GSH synthesis (QSH) to balance the ROS production. The hybrid “W/O” state is characterized by intermediate levels of both catabolic and anabolic activities. Finally, the “Q” state is highly reliant on glutamine oxidation. We will show that the dependence of glutamine oxidation is a robust feature of the “Q” state in the next section.

**Fig. 3:**
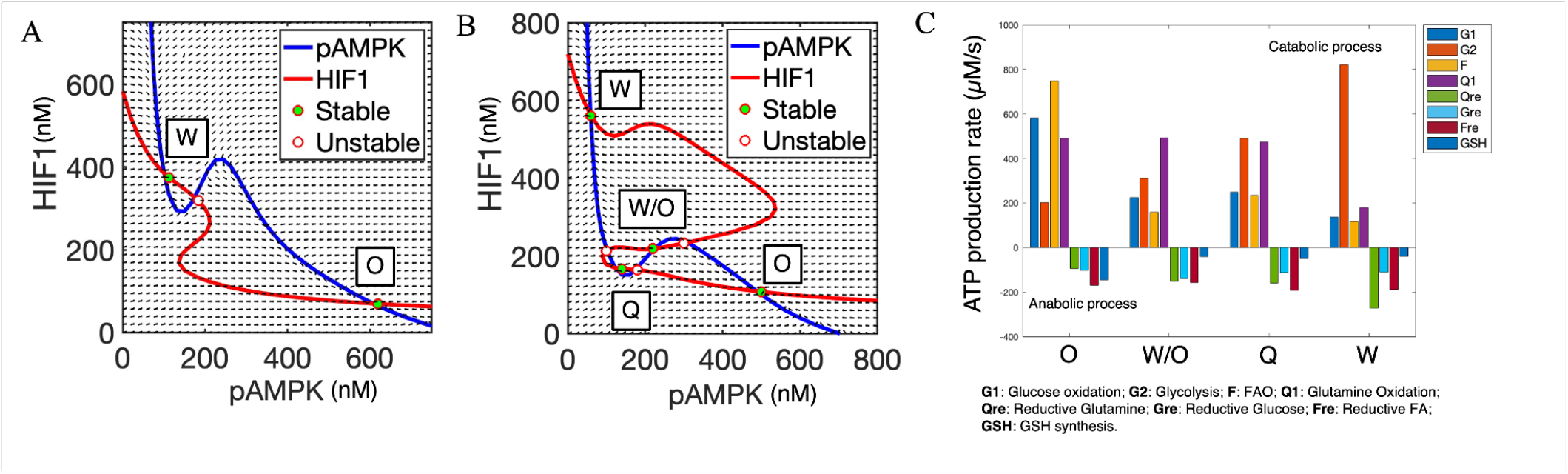
Model predicted association between gene states and metabolic pathway activities. **A**. Nullclines and steady states in the phase space of pAMPK and HIF-1 in normal cells. The red line represents the nullcline where the rate of change of HIF-1 is zero, and the blue line represents the nullcline where the rate of change of pAMPK is zero. Solid dots represent stable steady states while hollow dots represent unstable steady states. Each stable state is associated with a metabolic phenotype. Normal cells often acquire an OXPHOS phenotype, referred to as the state “O”, and a glycolytic phenotype when oxygen is limited, referred to as the state “W. **B**. The nullclines and steady states in the phase space of pAMPK and HIF-1 in cancer cells. Cancer cells can acquire two additional hybrid metabolic states – the ‘W/O’ state, characterized by intermediate HIF-1 and pAMPK activity and the ‘Q’ state, characterized by low pAMPK/HIF-1 but high glutamine oxidation. **C**. Net ATP production rates of different metabolic pathways for the “O”, “W/O”, “W”, and “Q” states. Positive rates represent catabolic processes where ATP is produced, while negative rates represent anabolic processes where ATP is consumed. Compared to the “W” state, the “O” state exhibits higher activities of glucose oxidation (G1), fatty acid oxidation (F), glutamine oxidation (Q1), and GSH synthesis (GSH), but lower activities of glycolysis (G2), reductive glutamine carboxylation (Qre), reductive glucose metabolism (Gre), and reductive fatty acid metabolism (Fre). The hybrid “W/O” state exhibits intermediate metabolic activities, while the “Q” state relies on high glutamine oxidation activity.

To identify the robustness of the stable states, we used a parameter randomization procedure. The overall strategy consists of randoming the model parameters for each simulation and collecting all stable state solutions from all simulations for statistical analysis to identify the robust solution patterns. For each Myc level, we considered 500 sets of model parameters. For each set of parameters, we randomly sampled from a uniform distribution of (75%*p*0, 125%*p*0), where *p*0 is the baseline value, and calculated the stable state solutions. Then we performed hierarchical clustering analysis of the results from all sets of parameters to identify the patterns of the solutions (**Fig. 4A**). We depicted the Myc level, metabolic state (which were defined as the overall activity of glucose oxidation/FAO/glycolysis to be compared with the previous study), glucose uptake and ATP production, ATP consumption, and the net values of ATP related to each cluster of solutions.

**Fig. 4.**
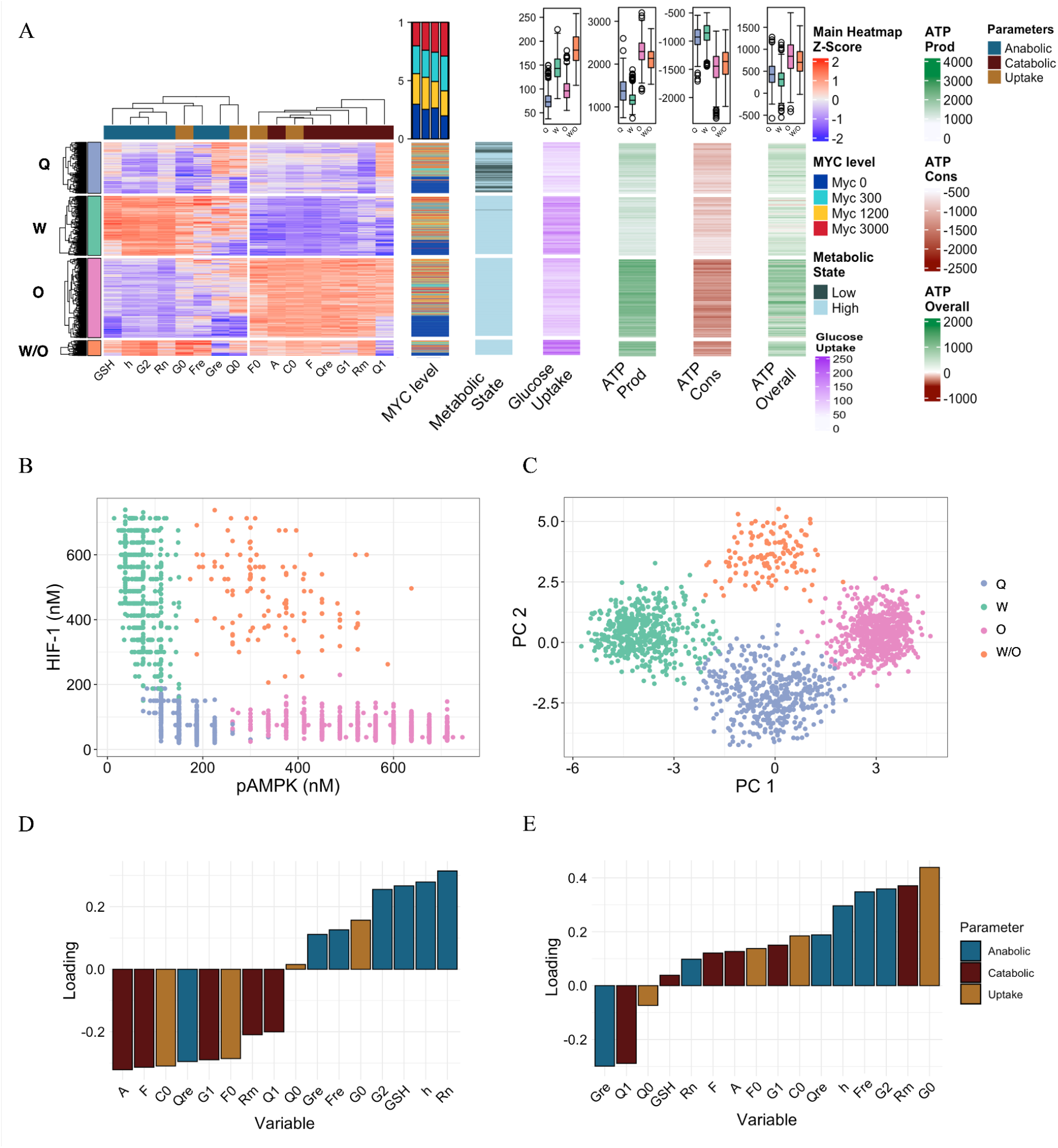
Hierarchical clustering analysis of stable state solutions. **A**. The primary heatmap displays four clusters of data derived from different Myc values (0, 300, 1200, 3000 nM). Each row in this heatmap represents a stable-state solution and each column represents one variable in the model. The proportions of each Myc group are shown in the first one-column heatmap, with a summary annotation at the top presenting the proportion of each Myc level within each cluster in a boxplot format (from left to right: “Q”, “W”, “O”, and “W/O”). The second one-column heatmap, labeled ‘Metabolic State’, corresponds to rows indicating whether steady states are active or inactive based on their glucose oxidation, glycolysis, and fatty acid oxidation values having Z-scores greater or less than 0, respectively. The third one-column heatmap shows the glucose uptake rates and the average for each cluster on the top, units expressed in µM/s. The subsequent one-column heatmaps, labeled ‘ATP Prod’, ‘ATP Cons’, and ‘ATP Overall’, illustrate the total ATP production, consumption, and net ATP for each row, taking into account both anabolic and catabolic processes. **B**. A plot of pAMPK vs HIF-1, illustrating the relationship between pAMPK and HIF-1 levels across all generated steady states. **C**. A scatter plot showing the results of a principal component analysis (PCA). Each point in the plot represents a steady state, plotted according to its scores on the first two principal components (PC1 and PC2) of the output element values in the minimal regulatory network. Different clusters are indicated by different colors, providing a visual representation of the grouping of steady states in the reduced-dimensional space of the PCA **D-E**. Bar plots showcasing the loading variables PC1(**D**) and PC2 (**E**) derived from the PCA. The variables are ranked based on their loading values, signifying their contribution to each principal component. Variable abbreviations: Glutathione Synthase (GSH), HIF-1 (h), Glycolysis (G2), NOx ROS (Rn), Glucose uptake (G0), Reductive Fatty acids (Fre), Reductive Glucose (Gre), Glutamine uptake (Q0), Fatty Acids uptake (F0), pAMPK (A), rate of acetyl-CoA for mitochondrial respiration (C0), FAO (F), Reductive Glutamine (Qre), Glucose oxidation (G1), mitochondrial ROS (Rm), glutamine oxidation (Q1).

By analyzing the stable state solutions from parameter randomization for each Myc value (0, 300nM, 1200nM, 3000nM representing an increasing level of Myc), four distinct groups of solutions were identified, corresponding to the four metabolic states - “W”, “O”, “W/O” and “Q” (**Fig. 4A**, left panel). The ‘O’ state and the ‘W’ state exhibit obviously opposite patterns of metabolic activities, as the ‘O’ state exhibits higher overall ATP production (as a result of higher ATP production and lower ATP consumption) relative to the ‘W’ state. Notably, the “W/O” state, which was previously defined as high AMPK/HIF-1/TCA/FAO/glycolysis, now has an improved characterization - high AMPK/HIF-1/catabolic/anabolic activities and exhibits the highest glucose uptake rates among the four metabolic phenotypes. The ‘Q’ state, which was previously defined as ‘low/low’ state with low AMPK/HIF-1/CTA/FAO/glycolysis, now has further characterization of high glutamine oxidation activity (Q1) and high reductive glucose activity (Gre). We confirmed that almost all the stable state solutions with negative Z-scores for glucose oxidation, glycolysis, and FAO, which were categorized as the ‘low/low’ state previously, were found to be in the “Q” states in the present study (**Fig. 4A**, second one-column heatmap).This indicates that cells in the previously defined ‘low/low’ state do not necessarily shut down all metabolic activities; instead, they tend to depend more on glutamine. This finding might have important implications for the use of metabolism-based therapy to overcome the problems of drug persistence. We also discovered that an increased level of Myc led to an increased proportion of the ‘Q’ states during the parameter randomization analysis, indicating an important role of Myc in the acquisition of the ‘Q’ state (**Fig. 4A**).

Next, we tested our conjecture that the two master gene regulators - pAMPK/HIF-1 activity can discriminate between these four metabolic states. By projecting all solutions on the pAMPK/HIF-1 axes, we showed a good distinction of the four states and a clear low pAMPK/HIF-1 signature of the ‘Q’ state (**Fig. 4B**). The difference between these metabolic states can also be visualized by principal component analysis (PCA) by projecting all the solutions along the first two principal components (PC1 and PC2) (**Fig. 4C**). PC1 mainly discriminates catabolic from anabolic variables, and discriminates between ‘W’, ‘O’, and (‘W/O’, ‘Q’) states (**Fig. 4D**). PC2 mainly discriminates between the ‘W/O’, (‘W’, ‘O’) and the ‘Q’ states (**Fig. 4E**). Furthermore, PC loadings indicate that if cells do not predominantly depend on glucose oxidation (G1) and FAO, they will compensate by resorting to glutamine oxidation (Q1). This metabolic shift enables cancer cells’ plasticity to meet their energy requirements and sustain their functions (21).

Our parameter randomization analysis shows that even upon relatively large perturbation to the model parameters, the characterization of the four metabolic phenotypes is robust. These results recapitulate many classical experimental observations, such as the ‘O’ state relying on OXPHOS and exhibiting high net value of ATP, with the ‘W’ state, with lower efficiency in ATP production, exhibiting increased glucose influx

### Model-predicted association between gene activity and metabolic pathway activity are confirmed in patient samples

To enable the testing of the model-predicted association between gene activity and metabolic pathway activity in each of the four metabolic states (“W”, “O”, “W/O”, and “Q”), we created scoring metrics for both gene regulators and metabolic pathways. To quantify metabolism, we applied our previously developed AMPK and HIF-1 signatures (14), glycolysis and FAO signatures (15), together with newly developed scoring metrics for Gre, Q1, GSH, Qre, Rn, Rm, Fre and Myc. The full list of genes for these metrics can be found on **SI Table 1**. For Fre and Myc, we followed the PCA-based method (14) to obtain the most relevant genes that contribute to the pathway activity. The scoring metric for metabolic pathways was determined by the mean expression of the Z-score of pertinent genes within each pathway. Then, we analyzed the RNA-seq data of patient samples of multiple cancer types from The Cancer Genome Atlas Program (TCGA).

We first applied hierarchical clustering analysis, taking into consideration all the genes of interest from the different anabolic and catabolic pathways (**SI Table 1**) in our model, to classify the patient samples into “W”, “W/O”, “O”, and “Q” states. Then we quantify the gene or metabolic pathway activity of the samples in each state. For example, based on the metabolic gene expression, the hepatocellular carcinoma (HCC) patient samples can be classified into four groups, corresponding to the “W”, “W/O”, “Q”, and “O” states (**Fig. 5A**). These four metabolic groups were also distinguishable according to their HIF-1 and pAMPK values: high HIF-1 and low pAMPK, intermediate HIF-1 and intermediate pAMPK, low HIF-1 and high pAMPK, and finally, low HIF-1 and low pAMPK (**Fig. S3A**). We include eleven one-column heat maps that illustrate the state of each evaluated metabolic parameter - (1) Myc activity, (2) HIF-1 activity, (3) AMPK activity, (4) glycolysis activity, (5) glucose oxidation, (6) FAO, (7) glutamine oxidation, (8) reductive glutamine metabolism, (9) overall catabolic activity and (11) overall anabolic activity. Next, we evaluated the predictions of our model pertaining to the unique characterizations of the four metabolic states. The model posits that the metabolic states “W” and “Q” demonstrate diminished glycolysis and fatty acid oxidation activities in comparison to the ‘O’ and ‘W/O’ states. This proposition aligns well with the analysis derived from the RNA-seq data of HCC samples (**Figs. 5B, S3A**). Furthermore, our model indicates that, in contrast to the ‘O’ and ‘W/O’ states, both the “W” and “Q” states display reduced catabolic activity. This is evidenced by the differences in ATP production (**Fig. 4A**). In the context of HCC, this difference is also discernible by the catabolic score, primarily characterized by decreased glucose oxidation and FAO. The ‘W’ state exhibits a higher anabolic activity relative to the ‘Q’ state. It is important to note that this anabolic state is characterized by glycolysis, HIF-1, and reductive activities (Fre, GSH, Qre) (**Fig. 4A**). This observation has been corroborated by the analysis of HCC patient samples (**Fig. 5B)**. Thus, our model does a good job of accounting for HCC data.

**Fig. 5.**
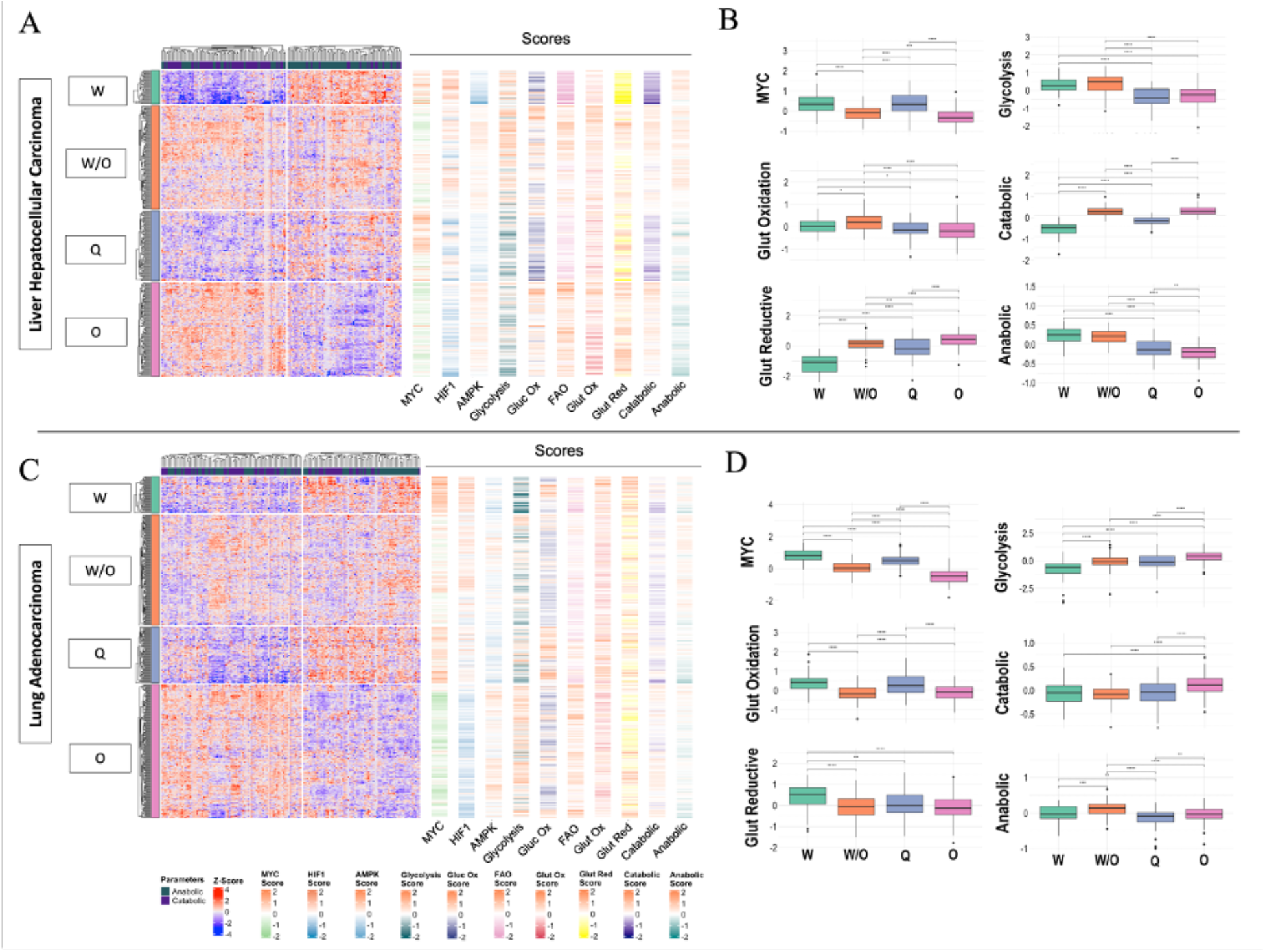
The association between gene activity and metabolic pathway activity. **A**. Heatmap of RNA-seq data from liver hepatocellular carcinoma (HCC). Each row represents a patient sample, and each column represents the expression of selected genes, which are divided according to their metabolic or anabolic activities. Four distinct clusters can be identified in the heatmap, corresponding to the four states identified in the metabolism model. The adjacent one-column heatmaps represent the scores for Myc, HIF-1, AMPK, glycolysis, glucose oxidation, FAO, glutamine oxidation, glutamine reduction, overall catabolic, and overall anabolic activities, respectively. **B**. Box plots summarizing and showing the differences of the scores according to each identified cluster in the HCC data. **C**. Heatmap of RNA-seq data from lung adenocarcinoma (LUAD). **D**. Box plots summarizing the differences in scores for each identified cluster in the LUAD data. For the box plots, a t-test was used to test the significance of each pair of clusters. Significance levels are indicated as follows: *, P < 0.05; **, P < 0.01; ***, P < 0.001.

Moving to the lung adenocarcinoma (LUAD) dataset, the LUAD samples also exhibit four distinct metabolic states - “W”, “O”, “W/O”, “Q” based on the metabolic gene expression (**Fig. 5D**) and can also be discriminated against based on their pAMPK and HIF-1 activity (**Figs. 5D-E, S3B**). The LUAD samples in the “W” and “Q” states present the expected low catabolic activity. The samples in the “O” state also align with our model-predicted characterization of high catabolic activity. For samples in the “W/O’’ state exhibit high anabolic and high catabolic activity. The results for many other cancer types can be found in the SI (**Fig. S4-10**). In general, samples in the “W/O’’ state exhibit both high anabolic and high catabolic activity. samples in the “Q” state exhibit high glutamine oxidation, except for the melanoma case (**Fig. S5**). Finally, for breast invasive carcinoma, tumors can be subclassified to different subtypes, namely luminal A, luminal B, HER2+, and basal-like. We show that the breast invasive carcinoma samples exhibit consistent characterization for the four metabolic states (**Fig. S9**). We further analyzed the relationship between the metabolic states and the cancer subtypes. Interestingly, we found that basal-like and HER2+ make up most of the “W” state that shows the highest Myc and high glucose consumption activity in comparison with samples in other metabolic states. All told, we confirmed the basic findings of our model-predicted metabolic characterization by analyzing the RNA-seq data of patient samples from TCGA.

### Quantification of glutamine metabolic activities

One novelty of the current metabolism model is the incorporation of glutamine metabolism and its master regulator - Myc. We next focus on studying the consumption of glutamine when cancer cells acquire different metabolic phenotypes. To relate to our previous characterization of the metabolic states (15), we performed a hierarchical clustering analysis of the stable state solutions based solely on the pAMPK and HIF-1 levels, and then determined the distribution of glutamine uptake (Q_0_) in each cluster (**Fig. 6A**). Our model suggests that in all four metabolic states, increased Myc level led to increased uptake of glutamine (**Fig. 6B**). To validate this model prediction, we assessed the predicted glutamine pathway activities of the “W”, “W/O”, “Q”, and “O” states. We applied a previously defined glutamine metabolism gene signature (GMGS, focusing on glutamine uptake) (22) and our defined Q_0_ signature (**Suppl Table 1**) to the breast cancer samples and 45 corresponding adjacent normal tissue samples for comparison. Notably, while the GMGS and the Q_0_ score share four genes, the Q_0_ signature includes additional genes that are involved in the transport of glutamine and derivatives (e.g. glutamate) into the mitochondria (e.g., SLC25A22, SLC25A13, SLC25A12). We showed that relative to normal cells, cancer cells exhibit much higher glutamine uptake rate (Q_0_) (**Fig. 6C bottom**, p <0.0001). . We found no significant difference between the GMGS signatures of tumor and normal samples (**Fig. 6C top**). We then segregated the 45 tumor samples based on our pAMPK/HIF-1 signatures into four groups. By both GMGS and our Q_0_ signature, we show that the samples in the “W” state exhibited significantly higher GMGS score relative to the samples in the “O” and “W/O” states. The result suggests that the “W” state has a more efficient uptake of glutamine (**Fig. 6D**). Interestingly, we showed that samples in the “Q” state also exhibit enhancements of glutamine uptake relative to the ‘O’ state (**Fig. 6D, bottom**).

**Figure 6:**
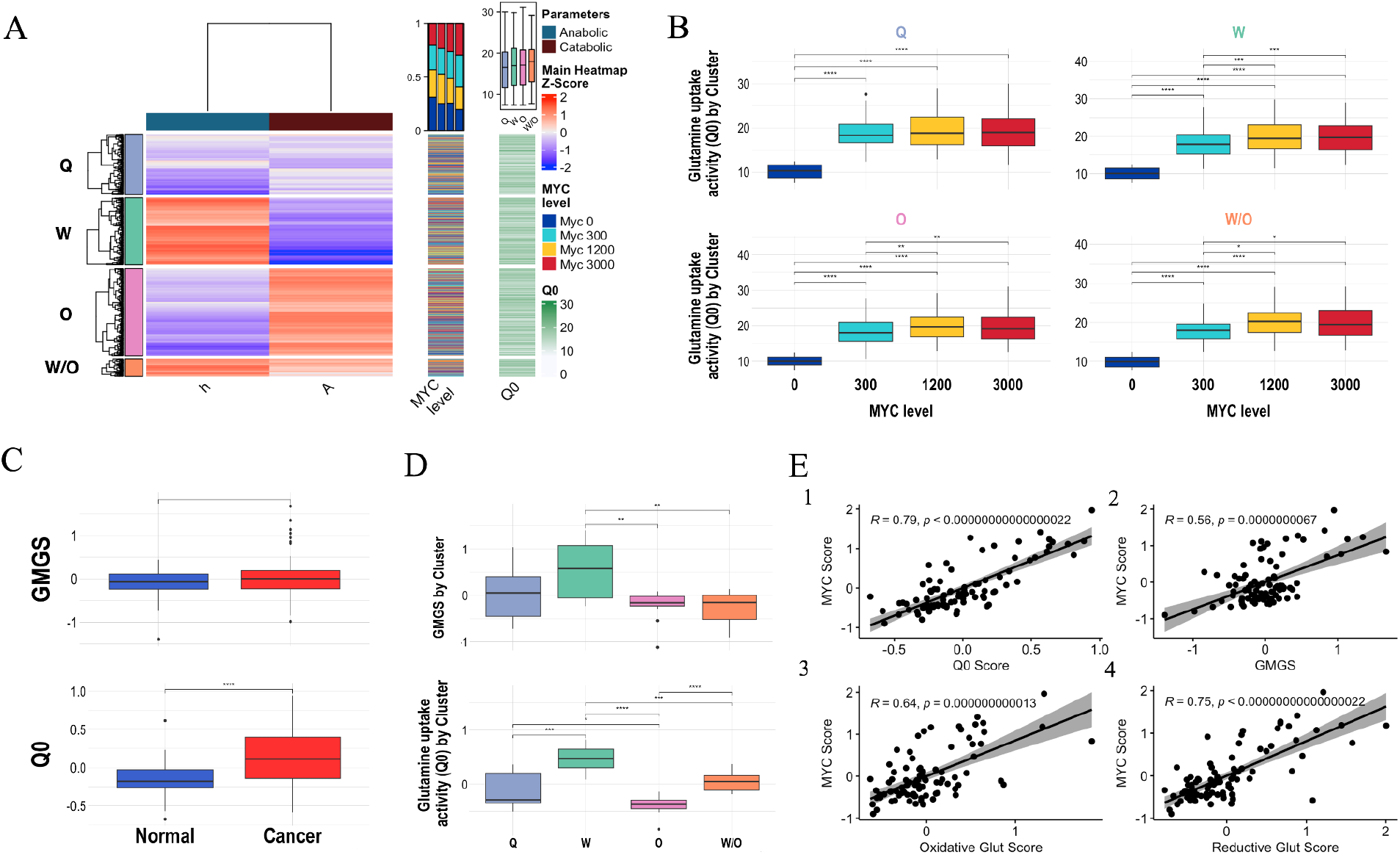
Model-predicted glutamine uptake and expression values for genes encoding glutamine-utilizing enzymes. **A**. Hierarchical clustering analysis of stable state solutions illustrating the relationship between the Glutamine uptake (Q0) levels and the states of pAMPK (A) and HIF-1 (h), the first one-column heat map indicates the Myc level (0, 300, 1200, 3000 nM) found in each state, the second one-column heat map shows the glutamine uptake activity for each identified state and the overall summarized at the top. **B**. Glutamine uptake identified in each cluster partitioned by Myc value. **C. Both** the previously published glutamine metabolism gene signature (GMGS) (22) and our glutamine uptake signature (Q0) were applied to a normal versus cancer dataset. **D**. Tumor samples were segregated into “W”, “O”, ‘W/O”, and “Q” groups to show their difference of the GMGS (top) and Q0 (bottom). **E**. Set of scatter plots comparing Q0 Score (1), GMGS (2), Oxidative Glut Score (3), and Reductive Glut Score (4), with the Myc Score. The regression line, confidence interval, and Pearson correlation coefficient are included. T-test was used to test the significance *, P < 0.05; **, P < 0.01; ***, P < 0.001.

As our model predicts a critical role of Myc in glutamine uptake rate in all metabolic phenotypes, we next evaluate how Myc activity is correlated with glutamine metabolic pathway activity. We found that there is a significantly positive correlation between the Myc score and the Q0 score (r = 0.78, p<0.001) (**Fig. 6E-1**), indicating that Myc significantly upregulates the glutamine uptake rate, aligning with our model expectation (**Fig. 6B**). Consistent results have been shown by the significant correlation between Myc and the GMGS signature (**Fig. 6E-2**) (r = 0.56, p<0.0001). Furthermore, we observed a strong correlation between Myc score and the glutamine oxidation score (**Fig. 6E-3**) and reductive glutamine metabolism score (**Fig. 6E-4**), suggesting an important role of Myc in different aspects of glutamine metabolism.

To quantify the activity of various glutamine-related metabolic pathways in the four metabolic states, we developed gene signatures for different metabolic pathways using involved genes. These genes can be categorized into *three* types - (1) enzyme genes involved in glutamine oxidation in mitochondria (e.g., GLS, GLS2, GOT2, GLUD1 and GPT2); (2) enzyme genes used in anabolic processes of glutamine (e.g., ASNS for asparagine synthesis, GFPT1 for hexosamine synthesis, PPAT for purine synthesis), and; (3) transporter genes that import glutamine (e.g., SLC38A1/2 and SLC1A5). Then we analyzed the RNA-seq data of liver hepatocellular carcinoma (HCC) and lung adenocarcinoma (LUAD) patient samples from The Cancer Genome Atlas Program (TCGA) to test the model-predicted characterization of glutamine metabolism in each of the four metabolic states (**Figs. S11-12**). After applying hierarchical clustering analysis and classifying the patient samples into “W”, “W/O”, “O”, and “Q” states, we quantified the expression of the glutamine metabolism genes among these four metabolic states. We found that for both HCC and LUAD samples, the samples in the state “O” and “W/O” states exhibit pronounced expression in most of the glutamine oxidation genes (GLS2, GOT2, and GPT2). The samples in the “W” or ‘W/O” state exhibit a pronounced GLS expression, while the samples in the “O” state exhibit a pronounced GLS2 expression. The inverse association between GLS and GLS2 expression has been reported before, and GLS is often over-expressed in cancer while GLS2 is regarded as a tumor suppressor (23). The tumor samples in group “W” and “Q” exhibit higher gene expression involved in amino acid/nucleotide synthesis (ASNS, GFPT1, and PPAT). The tumor samples in group “W” exhibit higher glutamine update (SLC38A1, SLC1A5) relative to samples in the “O”, “Q”, and “W/O” states, which is consistent with our findings by applying our Q0 or the GMGS signature (**Fig. 6D**). In summary, the “W” state exhibits high glutamine uptake and high anabolic processes involving glutamine, the “W/O” and the ‘O’ state exhibits high glutamine oxidation, whereas “Q” has genes high in glutamine oxidation and biosynthesis.

### The metabolic state of patient samples is significantly correlated with survival outcomes

Finally, we proceeded to examine the relationship between the metabolic state of patient samples with the overall survival. For the HCC patient samples (**Fig. 7A**), the ‘Q’ state was associated with the poorest survival outcome. Further inspection of the heatmap revealed that this particular cluster was characterized by elevated Myc activity and by high glutamine oxidation activity and high reductive glucose activity (**Figs. 5A-B**). The melanoma patient samples also exhibited the poorest survival outcome when associated with the “Q” state (**Fig. 7B**), Our previous study showed that the drug-tolerant melanoma cells exhibit low glucose/fatty acid metabolic activity, thus being characterized as the ‘low/low’ phenotype (16). All together, these drug-tolerant melanoma cells, which is associated with the ‘Q’ state probably rely on glutamine for survival. We further observed that colorectal adenocarcinoma, and leukemia exhibited the most unfavorable prognosis when in a hybrid ‘W/O’ (**Figs. 7C-D)**. This finding underscores the efficacy of our model in identifying metabolic states, especially the ‘W/O’ or ‘Q’ states that correlate with poor survival. For the LUAD dataset (**Fig. 7E**), the “O” state was associated with the best survival result, and both the LUAD dataset and the kidney cancer showed the worst outcome with the ‘W’ cluster having the highest Myc score value (Figs. 5C, 7E-F, S7B). Intriguingly, our analysis of the prostate cancer samples (**Fig. S13A**) revealed a distinct pattern. The most favorable prognosis was associated with the “Q” cluster. This particular cluster stands out from the rest of the cancer datasets due to a notable reduction in the Myc score (**Fig. S8**). This reduction appears to confer a survival advantage to the “Q” state, thereby underscoring the significant impact of Myc on survival outcomes. The rest of the cancers analyzed in this study, lung squamous cell carcinoma and breast invasive carcinoma, did not show any significant differences after being categorized by the four distinct phenotypes and analyzing the survival curves for each state (**Figs. S13B-C**). In short, there is some evidence that everything else being equal, hybrid states seem to be most aggressive. But, as we have seen for prostate cancer, other factors (such as Myc levels) are also critical and can overcome the purely metabolic effects.

**Fig. 7.**
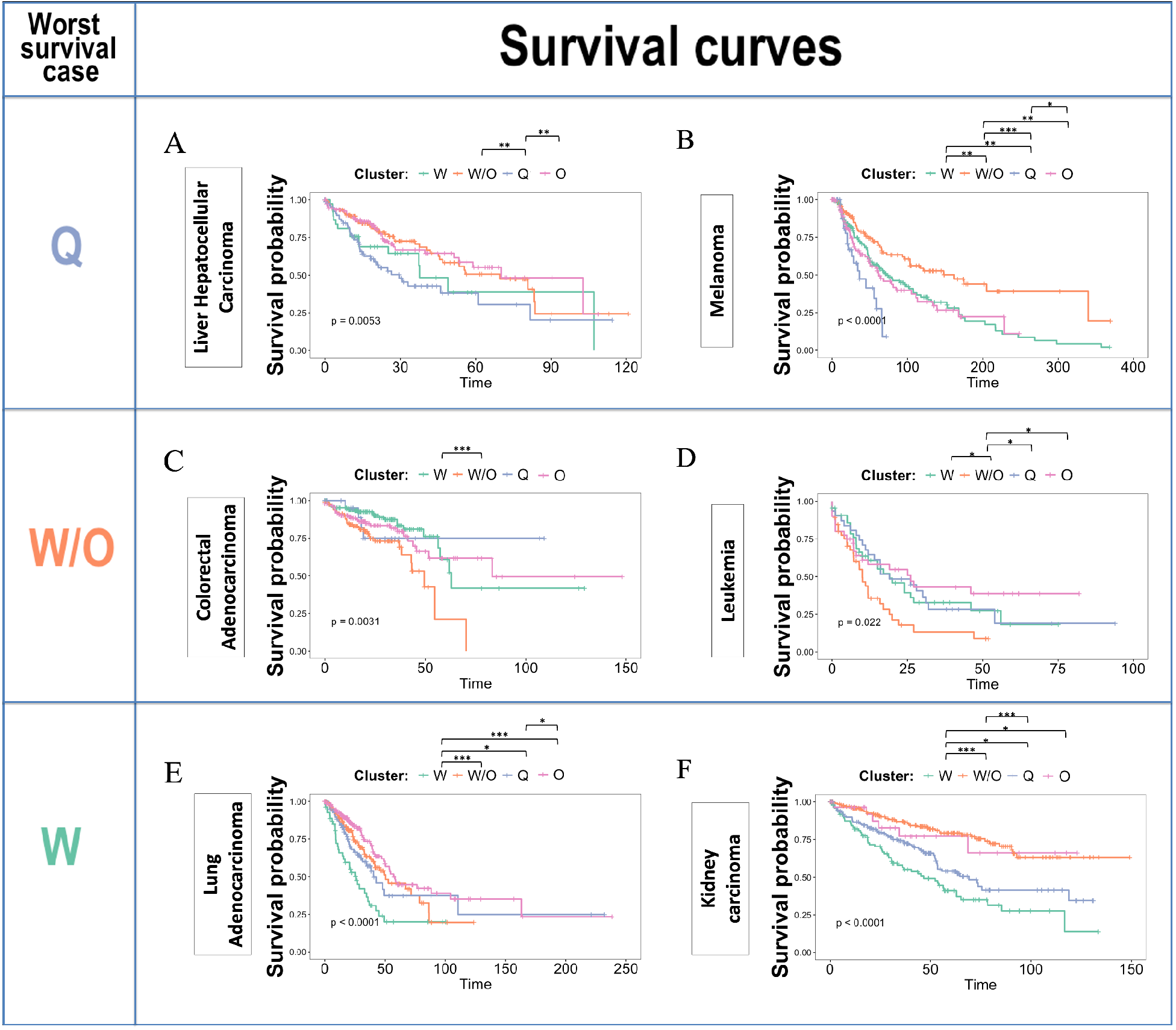
Survival curves stratified by metabolic states. Survival curves for liver hepatocellular carcinoma (A), melanoma (B), colorectal adenocarcinoma (C), leukemia (D), lung adenocarcinoma (E), and kidney carcinoma (F) datasets. Each dataset was separated into the four metabolic states – ‘O’, ‘W’, ‘W/O’ and ‘Q’. Each row in the figure indicates the metabolic state associated with the worst outcome for each cancer dataset. The survival curves were estimated using the Kaplan-Meier method and compared using the log-rank test. Pairwise comparisons of survival distributions stratified by metabolic state are displayed above the survival curve for each pair of metabolic state.Significance levels are indicated as follows: *, P < 0.05; **, P < 0.01; ***, P < 0.001.

## Discussion

Cancer cells demonstrate remarkable metabolic versatility by utilizing a variety of nutrients such as glucose, fatty acids, and glutamine for both catabolic and anabolic processes. These metabolic pathways not only generate ATP but also produce intermediates essential for biosynthesis, supporting rapid cell proliferation. This versatility of cancer metabolism enables cancer cells to thrive in diverse microenvironments and contributes to their resistance against therapeutic interventions (24). To shed light on cancer metabolic versatility, we have developed a comprehensive mathematical model that, for the first time, integrates three indispensable metabolic ingredients (glucose, fatty acids, and glutamine) and captures the complex interplay of nutrient uptake and utilization and the dynamic shifts between energy production and biomolecule synthesis.

Through modeling, we demonstrated that cancer cells can achieve four main metabolic states, W, O, W/O and Q, characterized by both gene activity (AMPK/HIF/Myc) and metabolic pathway activity. We show that the ‘O’ state, that was characterized by high AMPK and high OXPHOS, is high in overall catabolic activity, while the ‘W’ state, that was characterized by high HIF-1 and high glycolysis, is high in overall anabolic activity. We provide finer resolution for the hybrid W/O state and the low/low state. We show that the hybrid ‘W/O’ state exhibits both high catabolic (glucose oxidation, fatty acid oxidation not glutamine oxidation) and high anabolic activities (reductive fatty acid and reductive glutamine but not glucose). We show that the previously defined ‘low/low’ state, while exhibiting low overall catabolic/anabolic activity relative to the rest states, exhibits pronounced glutamine oxidation activity (therefore we refer to this state as ‘Q0’ in this manuscript) and reductive glucose metabolism. Our model recapitulates the critical role of Myc in glutamine metabolism. The model suggests higher levels of Myc upregulates glutamine uptake in all four metabolic states and promotes both glutamine oxidation and reductive glutamine metabolism.

These predicted metabolic characterizations have been confirmed by analyzing RNA-seq data in patient samples from TCGA. Moreover, the model delineated the association between gene activity and metabolic pathway activity, which has also been verified by analyzing the patient sample data. Furthermore, we evaluated the functional consequences of different metabolic states of patient samples. We observed that patient samples characterized as metabolic hybrids exhibit the worst overall survival outcomes relative to samples with other metabolic states in hepatocellular cancer, colorectal adenocarcinoma, melanoma, and leukemia. One limitation of the current study is that Myc was treated as an input to the network. In the future, integrating the detailed feedback from other gene regulators and metabolic intermediates to Myc would improve our understanding of the effect of Myc. In summary, our integrated modeling-data analysis approach provides a holistic understanding of cancer metabolism and an extendable framework for including additional biological factors.

A promising direction would be to enhance the current metabolism model by coupling it with other biological processes that consume ATP and/or biomass in cancer cells. This could be done by determining the parameters in the current model as functions of additional processes, such as cell migration, and division. This would be an improvement on the current modular modeling of the cancer process. For instance, the epithelial-mesenchymal transition (EMT) influences how cells acquire migratory and invasive properties. By incorporating EMT into our metabolism model, we can simulate how EMT affects nutrient utilization, and vice versa, how changes in metabolism affects EMT (25). This would help identify critical points of intervention where targeting metabolic adaptations and EMT dynamics concurrently may yield synergistic therapeutic benefits. Another important unresolved question is how cell metabolism regulates tumor dormancy, which places patients at risk of metastatic relapse for the remainder of their life. It has been observed that whether tumor cells are proliferative or quiescent can result from a ‘tug of war’ between oxidative stress and antioxidative response. Coupling the metabolism model with the molecular networks regulating tumor dormancy processes should yield significant insights (26). One speculation is there may be a connection between the “Q” state which exhibits low activities of many metabolic processes (aside from glutamine oxidation) and tumor dormancy. Another promising direction is to extend the metabolism model from intracellular level to intercellular cell by considering the competition for resources between cancer cells and immune cells (27). Altogether, our metabolism model serves as a valuable tool for identifying potential metabolic vulnerabilities and designing targeted interventions to effectively disrupt metabolism relevant processes in cancer cells.

## Acknowledgments

We acknowledge useful conversations with Benny Kaipperettu at the outset of this project. HL and JNO acknowledge NSF support of the Center for Theoretical Biological Physics, PHY-2019745. HL was also supported in part by NSF grant DMS-2245957. JNO was also supported by the NSF grant PHY-2210291. JVC acknowledges CONAHCYT for financial support granted for PhD Studies (CVU 637952). JTG was supported by the Cancer Prevention and Research Institute of Texas (CPRIT RR210080). JTG is a CPRIT Scholar in Cancer Research.

