## Supplementary Information for "Computational modeling of cancer cell metabolism along the catabolic-anabolic axes"

**Methods:**

Our model consists of the following coupled equations:

$$\dot{R}_{mt} = \left(\gamma_{G_1} * G_1 + \gamma_F * F + \gamma_{Q_1} * Q_1\right) - k_{R_{mt}} * R_{mt} * H^{s+}\left(A, A^0_{R_{mt}}, \lambda_{A,R_{mt}}, n_{A,R_{mt}}\right) * H^{s+}\left( G_{GSH}, G^0_{GSH,R}, \lambda_{GSH,R}, n_{GSH,R}\right) \quad (\boldsymbol{eq.1})$$

$$\dot{R}_{nox} = g_{R_{nox}} \cdot C^{comp}_{R_{nox}}\left(g_0, H, g_{H,R_{nox}}, H^0_{R_{nox}}, n_{H,R_{nox}}, A, g_{A,R_{nox}}, A^0_{R_{nox}}, n_{A,R_{nox}}\right) - k_{R_{nox}} * R_{nox} * H^{s+}\left( G_{GSH}, G^0_{GSH,R}, \lambda_{GSH,R}, n_{GSH,R}\right) \quad (\boldsymbol{eq.2})$$

$$R_T = R_{mt} + R_{nox} \quad (\boldsymbol{eq.3})$$

$$\dot{A} = g_A * H^{s-}\left(H, H^0_A, \lambda_{H,A}, n_{H,A}\right) * H^{s+}\left(R_T, R^0_{T,A}, \lambda_{R_T,A}, n_{R_T,A}\right) * H^{s-}\left( X_{ATP}, X^0_{ATP,A}, \lambda_{X_{ATP},A}, n_{X_{ATP},A}\right) - k_A * A \quad (\boldsymbol{eq.4})$$

$$\dot{H} = g_H * H^{s-}\left(A, A^0_H, \lambda_{A,H}, n_{A,H}\right) - k_H * H * H^{s-}\left(G_2, G^0_{2,H}, \lambda_{G_2,H}, n_{G_2,H}\right) * H^{s-}\left(R_T, R^0_{T,H}, \lambda_{R_T,H}, n_{R_T,H}\right) * \left(M, M^0_{M,H}, \lambda_{M,H}, n_{M,H}\right) \quad (\boldsymbol{eq.5})$$

$$G_0 = g_{G_0} * [H^{S+}\left(H, H^0_{G_0}, \lambda_{H,G_0}, n_{H,G_0}\right) + H^{S+}\left(A, A^0_{G_0}, \lambda_{A,G_0}, n_{A,G_0}\right) + H^{S+}\left(M, M^0_{G_0}, \lambda_{M,G_0}, n_{M,G_0}\right)] \quad (\boldsymbol{eq.6})$$

$$Q_0 = g_{Q_0} * H^{s+}\left(M, M^0_{Q_0}, \lambda_{M,Q_0}, n_{M,Q_0}\right) \quad (\boldsymbol{eq.7})$$

$$F_0 = g_{F_0} * H^{s+}\left(A, A^0_{F_0}, \lambda_{A,F_0}, n_{A,F_0}\right) \quad (\boldsymbol{eq.8})$$

$$C_0 = g_{C_0} * H^{s+}\left(A, A^0_{C_0}, \lambda_{A,C_0}, n_{A,C_0}\right) \quad (\boldsymbol{eq.9})$$

$$G = G_1 + G_2 + G_r \quad (\boldsymbol{eq.10})$$

$$Q = Q_1 + Q_r + Q_{GSH} \quad (\boldsymbol{eq.11})$$

$$F = F_1 + F_r - \gamma_{G,F} * G_r - \gamma_{G,Q} * Q_r \quad (\boldsymbol{eq.12})$$

$$C = 2 * G_1 + 9 * F \quad (\boldsymbol{eq.13})$$

$$G_1 = g_{G_1} * H^{s-}\left(G, G_0, \lambda_{G,G_1}, n_{G,G_1}\right) * H^{s-}\left(C, C_0, \lambda_{C,G_1}, n_{C,G_1}\right) \quad (\boldsymbol{eq.14})$$

$$G_2 = g_{G_2} * H^{s-}\left(G, G_0, \lambda_{G,G_2}, n_{G,G_2}\right) * H^{s+}\left(H, H^0_{G_2}, \lambda_{H,G_2}, n_{H,G_2}\right) \quad (\boldsymbol{eq.15})$$

$$G_r = g_{G_r} * H^{s-}\left(G, G_0, \lambda_{G,G_{re}}, n_{G,G_{re}}\right) * H^{s-}\left(A, A^0_{G_{re}}, \lambda_{A,G_{re}}, n_{A,G_{re}}\right) * H^{s+}\left(M, M^0_{G_{re}}, \lambda_{M,G_{re}}, n_{M,G_{re}}\right) \quad (\boldsymbol{eq.16})$$

$$F_1 = g_{F_1} * H^{s-}(F, F_0, \lambda_{F,F_1}, n_{F,F_1}) * H^{s-}(C, C_0, \lambda_{C,F}, n_{C,F}) * H^{s+}(A, A_F^0, \lambda_{A,F}, n_{A,F}) * H^{s-}(H, H_F^0, \lambda_{H,F}, n_{H,F}) \quad (\boldsymbol{eq.17})$$

$$F_r = g_{F_r} * H^{s-}(F, F_0, \lambda_{F,F_r}, n_{F,F_r}) \ (\boldsymbol{eq.18})$$

$$Q_1 = g_{Q_1} \cdot H^{s-}(Q, Q_0, \lambda_{Q,Q_1}, n_{Q,Q_1}) \cdot H^{s+}(M, M_{Q_1}^0, \lambda_{M,Q_1}, n_{M,Q_1}) \cdot H^{s-}(H, H_{Q_1}^0, \lambda_{H,Q_1}, n_{H,Q_1}) \quad (\boldsymbol{eq.19})$$

$$Q_{GSH} = g_{Q_{GSH}} \cdot H^{s-}(Q, Q_0, \lambda_{Q,Q_{GSH}}, n_{Q,Q_{GSH}}) \cdot H^{s-}(M, M_{Q_{GSH}}^0, \lambda_{M,Q_{GSH}}, n_{M,Q_{GSH}}) \quad (\boldsymbol{eq.20})$$

$$\dot{G_{GSH}} = Q_{GSH} - k_{GSH} * G_{GSH} \ (\boldsymbol{eq.21})$$

$$Q_r = g_{Q_r} \cdot H^{s-}(Q, Q_0, \lambda_{Q,Q_r}, n_{Q,Q_r}) \cdot H^{s+}(M, M_{Q_r}^0, \lambda_{M,Q_r}, n_{M,Q_r}) \cdot H^{s+}(H, H_{Q_r}^0, \lambda_{H,Q_r}, n_{H,Q_r}) \cdot H^{s-}(A, A_{Q_r}^0, \lambda_{A,Q_r}, n_{A,Q_r}) \quad (\boldsymbol{eq.22})$$

$$G_{1,ATP} = 29 * G_1 \quad (\boldsymbol{eq.23})$$

$$G_{2,ATP} = 2 * G_2 \quad (\boldsymbol{eq.24})$$

$$Q_{1,ATP} = 24 * Q_1 \ (\boldsymbol{eq.25})$$

$$F_{1,ATP} = 106 * F_1 \ (\boldsymbol{eq.26})$$

$$F_{2,ATP} = 7 * F_2 \ (\boldsymbol{eq.27})$$

$$G_{r,ATP} = (15 - 2) * G_r \ (\boldsymbol{eq.28})$$

$$Q_{r,ATP} = 15 * Q_r \ (\boldsymbol{eq.29})$$

$$Q_{GSH,ATP} = 2 * Q_{GSH} \ (\boldsymbol{eq.30})$$

$$X_{ATP} = G_{1,ATP} + G_{2,ATP} + Q_{1,ATP} + F_{1,ATP} - G_{r,ATP} - Q_{r,ATP} - Q_{GSH,ATP} \ (\boldsymbol{eq.31})$$

We now explain these in detail. To simulate the temporal dynamics of the regulatory proteins pAMPK and HIF-1 as well as the temporal dynamics of metabolites mtROS and noxROS, we devise the following equations ($eqs. 1 - 5$).

$$\dot{R}_{mt} = g_{R_{mt}} \cdot (\gamma_{G_1} G_1 + \gamma_F F + \gamma_{Q_1} Q_1) - k_{R_{mt}} \cdot R_{mt} \cdot (H^{s+}(A, A_{R_{mt}}^0, \lambda_{A,R_{mt}}, n_{A,R_{mt}}) + \gamma_{GSH} Q_{GSH}) \quad (\boldsymbol{eq.1})$$

($eq. 1$) **represents the temporal dynamics of mitochondrial reactive oxygen species (mtROS) ($R_{mt}$).** $g_{R_{mt}}$ is the basal production rate of mtROS, $(\gamma_{G_1} G_1 + \gamma_F F + \gamma_{Q_1} Q_1)$ represents the increase of mtROS production due to glucose oxidation ($G_1$), FAO ($F$) and glutamine oxidation ($Q_1$). Notably, the two parameters $\gamma_{G1}$ and $\gamma_F$ have fixed ratio 2/9 because the ratio of the amount of acetyl-CoA entering TCA generated by glucose oxidation and FAO is 2/9. $k_{R_{mt}}$ represents the basal degradation rate of mtROS and the shifted Hill function (1) $H^{s+}(A\ , A^0_{R_{mt}}, \lambda_{A,R_{mt}}, n_{A,R_{mt}})$ represents the deoxidation effect of AMPK. $\gamma_{GSH}G_{GSH}$ represents the antioxidation effect of glutathione (GSH) due to the GSH synthesis pathway ($Q_{GSH}$).

$$\dot{R}_{nox} = g_{R_{nox}} \cdot C^{comp}_{R_{nox}}(g_0, H, g_{H,R_{nox}}, H^0_{R_{nox}}, n_{H,R_{nox}}, A, g_{A,R_{nox}}, A^0_{R_{nox}}, n_{A,R_{nox}}) \\ -k_{R_{nox}} \cdot R_{nox} \cdot (1 + \gamma_{GSH}Q_{GSH}) \quad (\boldsymbol{eq.2})$$

($eq.2$) **represents the temporal dynamics of NADPH Oxidase-derived Reactive Oxygen Species (noxROS) ($R_{nox}$).** $g_{R_{nox}}$ is the basal production rate of noxROS, $C^{comp}_{R_{nox}}(g_0, H, H^0_{HR_{nox}}, n_{H_{nox}}, g_1, A, A_0, g_2, n_{A_{nox}})$ represents the competitive regulation of noxROS production by AMPK ($A$) and HIF-1 ($H$) and $k_{R_{nox}}$ represents the basal degradation rate of noxROS. $\gamma_{GSH}G_{GSH}$ represents the antioxidation effect of GSH due to the GSH synthesis pathway ($G_{GSH}$).

$$R_T = R_{mt} + R_{nox} \quad (\boldsymbol{eq.3})$$

($eq.3$) **represents the total level of ROS ($R_T$), which is the sum of mtROS ($R_{mt}$) and noxROS ($R_{nox}$).**

$$\dot{A} = g_A \cdot H^{s+}(R_T, R^0_{T,A}, \lambda_{R_T,A}, n_{R_T,A}) \cdot H^{s-}(H, H^0_A, \lambda_{H,A}, n_{H,A}) \cdot H^{s-}(X_{ATP}, X^0_{ATP,A}, \lambda_{X_{ATP},A}, n_{X_{ATP},A}) \\ - k_A A \quad (\boldsymbol{eq.4})$$

($eq.4$) **represents the temporal dynamics of phosphorylated AMPK (pAMPK) ($A$).** $g_A$ is the basal production rate of pAMPK, $H^{s+}(R_T\ , R^0_{T,A}, \lambda_{R_T,A}, n_{R_T,A})$ represents the excitatory regulation on pAMPK production by ROS ($R_T$), $H^{s-}(H\ , H^0_{H,A}, \lambda_{H,A}, n_{H,A})$ represents the inhibitory regulation on pAMPK by HIF-1 ($H$), $H^{s-}(X_{ATP}\ , X^0_{ATP,A}, \lambda_{X_{ATP},A}, n_{X_{ATP},A})$ represents the inhibitory regulation on pAMPK by ATP ($X_{ATP}$) and $k_A$ represents the basal degradation rate of AMPK.

$$\dot{H} = g_H \cdot H^{s-}(A, A^0_H, \lambda_{A,H}, n_{A,H}) - k_H \cdot H \cdot H^{s-}(G_2, G^0_{2,H}\ , \lambda_{G_2,H}, n_{G_2,H}) \\ \cdot H^{s-}(R_T, R^0_{T,H}, \lambda_{R_T,H}, n_{R_T,H}) \cdot H^{s-}(M, M^0_{M,H}, \lambda_{M,H}, n_{M,H}) \quad (\boldsymbol{eq.5})$$

($eq.5$) **represents the temporal dynamics of HIF-1 ($H$).** $g_H$ is the basal production rate of HIF-1, $H^{s-}(A\ , A^0_H, \lambda_{A,H}, n_{A,H})$ represents the inhibitory regulation on HIF-1 production by pAMPK, $k_H$ represents the basal degradation rate of HIF-1, the shifted Hill functions $H^{s-}(G_2\ , G^0_{2,H}\ , \lambda_{G_2,H}, n_{G_2,H})$, $H^{s-}(R_T\ , R^0_{T,H}, \lambda_{R_T,H}, n_{R_T,H})$ and $H^{s-}(M, M^0_{M,H}, \lambda_{M,H}, n_{M,H})$ represent the stabilization of HIF-1 by the glycolytic activity ($G_2$), ROS ($R_T$) and Myc ($M$).

Since the chemical reactions in the metabolism processes are much faster than the genetic regulations, we assume the metabolites and the pathways are in the equilibrium state at certain level of pAMPK and HIF-1. To capture the dynamics of metabolic flux, we derive the following equations ($eqs. 6 - 16$).

$$G_0 = g_{H,G_0} \cdot H^{s+}(H, H^0_{G_0}, \lambda_{H,G_0}, n_{H,G_0}) + g_{A,G_0} \cdot H^{s+}(A, A^0_{G_0}, \lambda_{A,G_0}, n_{A,G_0}) \\ + g_{M,G_0} \cdot H^{s+}(M, M^0_{G_0}, \lambda_{M,G_0}, n_{M,G_0}) \quad (\boldsymbol{eq.6})$$

$(eq.6)$ **represents the glucose uptake rate** $(G_0)$**.** Since HIF-1, pAMPK and Myc can enhance the glucose uptake, one assumption is the maximum glucose uptake rate $(G_0)$ is determined by the HIF-1, pAMPK and Myc levels. $g_{H,G_0}H^{s+}(H, H_{G_0}^0, \lambda_{H,G_0}, n_{H,G_0})$ represents the regulation of glucose uptake by HIF-1, $g_{A,G_0}H^{s+}(A, A_{G_0}^0, \lambda_{A,G_0}, n_{A,G_0})$ represents the regulation of glucose uptake by pAMPK and $g_{M,G_0}H^{s+}(M, M_{G_0}^0, \lambda_{M,G_0}, n_{M,G_0})$ represents the regulation of glucose uptake by Myc.

$$Q_0 = g_{M,G_{Gln,0}} \cdot H^{s+}(M, M_{Q_0}^0, \lambda_{M,Q_0}, n_{M,Q_0}) \quad (\boldsymbol{eq.7})$$

$(eq.7)$ **represents the glutamine uptake rate (**$G_{Gln,0}$**).** As Myc can enhance glutamine update, one assumption is the maximum glutamine uptake rate $(G_{Gln,0})$ is determined by the Myc levels.

$$F_0 = g_{F_0} \; * H^{s+}(A, A_{F_0}^0, \lambda_{A,F_0}, n_{A,F_0}) \quad (\boldsymbol{eq.8})$$

$(eq.8)$ **represents the fatty acid uptake rate** $(F_0)$**.**

$$C_0 = g_{A,C_0} \cdot H^{s+}(A, A_{C_0}^0, \lambda_{A,C_0}, n_{A,C_0}) \quad (\boldsymbol{eq.9})$$

$(eq.9)$ **represents the maximum utilization rate of acetyl-CoA for mitochondrial respiration** $(C_0)$**.** Since the rate of acetyl-CoA entering the TCA cycle is limited by the mitochondrial activity that is determined by the pAMPK, one assumption here is the maximum utilization rate of Acetyl-CoA $(C_0)$ is determined by the pAMPK levels. The shifted Hill function $g_{A,C_0}H^{s+}(A\ , A_{C_0}^0, \lambda_{A,C_0}, n_{A,C_0})$ represents the regulation of pAMPK on the utilization of Acetyl-CoA for mitochondrial TCA cycle.

The glucose uptake rate $(G_0)$ and the utilization rate of Acetyl-CoA $(C_0)$ for TCA cycle restrict the activities of three metabolic pathway – glucose oxidation $(G_1)$, glycolysis $(G_2)$, and FAO $(F)$.

$$G = G_1 + G_2 + G_{re} \quad (\boldsymbol{eq.10})$$

$(eq.10)$ **represents the glucose consumption rate** $(G)$**.** $G_1$ represents the total glucose consumption rate, which is equal to the sum of the glucose oxidation rate $(G_1)$ and glycolysis rate $(G_2)$ and the reductive glucose metabolic rate $(G_{re})$ (mainly representing fatty acid synthesis rate from glucose), since glucose is shared by these pathways.

$$Q = Q_1 + Q_{GSH} + Q_{re} \quad (\boldsymbol{eq.11})$$

$(eq.11)$ **represents the total glutamine consumption rate (**$Q$**),** which is the sum of glutamine oxidation rate $(Q_1)$ and glutathione (GSH) synthesis rate $(Q_{GSH})$ and the reductive glutamine metabolic rate $(Q_{re})$ (mainly representing fatty acid synthesis rate from glutamine), since intracellular glutamine is shared by these three pathways.

$$F = F_1 + F_r - \gamma_{G,F} * G_r - \gamma_{G,Q} * Q_r \; (\boldsymbol{eq.12})$$

$(eq.12)$ **represents the total fatty acid consumption rate (**$F$**),**

$$C = 2 * G_1 + 9 * F \quad (\boldsymbol{eq.13})$$

$(eq. 13)$ **represents the production rate of Acetyl-CoA for mitochondrial respiration** $(C)$**.** The generated Acetyl-CoA that can enter the TCA cycle for ATP production is determined by glucose oxidation rate $(G_1)$ and FAO rate $(F)$. 2 molecules of acetyl-CoA is produced in 1 glucose oxidation process, and 9 molecules of acetyl-CoA is produced in 1 FAO process, in which we assumed the average carbon atoms contained in each fatty acid is 18.

$$G_1 = g_{G_1} H^{s-}(G, G_0, \lambda_{G,G_1}, n_{G,G_1}) H^{s-}(C, C_0, \lambda_{C,G_1}, n_{C,G_1}) \quad (\boldsymbol{eq. 14})$$

$$G_2 = g_{G_2} H^{s-}(G, G_0, \lambda_{G,G_2}, n_{G,G_2}) H^{s+}(H, H^0_{G_2}, \lambda_{H,G_2}, n_{H,G_2}) \quad (\boldsymbol{eq. 15})$$

$$G_r = g_{G_r} \cdot H^{s-}(G, G_0, \lambda_{G,G_{re}}, n_{G,G_{re}}) \cdot H^{s-}(A, A^0_{Gre}, \lambda_{A,G_{re}}, n_{A,G_{re}}) \cdot H^{s+}(M, M^0_{Gre}, \lambda_{M,G_{re}}, n_{M,G_{re}}) (\boldsymbol{eq. 16})$$

$$F_1 = g_{F_1} * H^{s-}(F, F_0, \lambda_{F,F_1}, n_{F,F_1}) * H^{s-}(C, C_0, \lambda_{C,F}, n_{C,F}) * H^{s+}(A, A^0_F, \lambda_{A,F}, n_{A,F}) * H^{s-}(H, H^0_F, \lambda_{H,F}, n_{H,F}) \quad (\boldsymbol{eq. 17})$$

$$F_r = g_{F_r} * H^{s-}(F, F_0, \lambda_{F,F_r}, n_{F,F_r}) \; (\boldsymbol{eq. 18})$$

$(eqs. 12 - 18)$ **represent the glucose oxidation rate** $(G_1)$**, the glycolysis rate** $(G_2)$**, the reductive glucose metabolic rate** $(G_r)$**, the FAO rate** $(F_1)$**, and the reductive fatty acid metabolism** $(F_r)$ **respectively.**

The negative shifted Hill functions $H^{s-}(G, G_0, \lambda_{G,G_1}, n_{G,G_1})$, $H^{s-}(G, G_0, \lambda_{G,G_2}, n_{G,G_2})$ and $H^{s-}(G, G_0, \lambda_{G,G_{re}}, n_{G,G_{re}})$ represent the competition of glucose oxidation $(G_1)$, glycolysis $(G_2)$ and reductive glucose metabolism $(G_{re})$ on glucose utilization. The threshold $G_0$, that is the glucose uptake rate, in these three shifted Hill functions adds restriction on $G_1$, $G_2$ and $G_{re}$ since if $G > G_0$, that means the glucose utilization rate is larger than the glucose uptake rate, these negative shifted Hill functions will decrease $G_1$, $G_2$ and $G_{re}$ thus decreasing $G$.

The negative shifted Hill functions $H^{s-}(C, C_0, \lambda_{C,G_1}, n_{C,G_1})$ and $H^{s-}(C, C_0, \lambda_{C,F}, n_{C,F})$ representing the competition of glucose oxidation $(G_1)$ and FAO $(F)$ on acetyl-CoA production. The threshold $C_0$, that is the limiting utilization rate of acetyl-CoA for mitochondrial respiration, in these two shifted Hill functions adds restriction on both $G_1$ and $F$ since if $C > C_0$, that means the produced acetyl-CoA is beyond the limiting utilization rate of acetyl-CoA for mitochondrial respiration, these negative shifted Hill functions will decrease both $G_1$ and $F$ thus decreasing $C$.

$H^{s+}(H, H^0_{G_2}, \lambda_{H,G_2}, n_{H,G_2})$ in $(eq. 11)$ represents the regulation of glycolytic activity by HIF-1.
$H^{s+}(A, A^0_F, \lambda_{A,F}, n_{A,F})$ in $(eq. 12)$ represents the regulation of FAO by AMPK.
$H^{s-}(A, A^0_{Gre}, \lambda_{A,G_{re}}, n_{A,G_{re}})$ represents the inhibition of fatty acid synthesis by AMPK.

$$Q_1 = g_{Q_1} \cdot H^{s-}(Q, Q_0, \lambda_{Q,Q_1}, n_{Q,Q_1}) \cdot H^{s+}(M, M^0_{Q_1}, \lambda_{M,Q_1}, n_{M,Q_1}) \cdot H^{s-}(H, H^0_{Q_1}, \lambda_{H,Q_1}, n_{H,Q_1}) \quad (\boldsymbol{eq. 19})$$

$$Q_{GSH} = g_{Q_{GSH}} \cdot H^{s-}(Q, Q_0, \lambda_{Q,Q_{GSH}}, n_{Q,Q_{GSH}}) \cdot H^{s-}(M, M^0_{Q_{GSH}}, \lambda_{M,Q_{GSH}}, n_{M,Q_{GSH}}) \quad (\boldsymbol{eq. 20})$$

$$\dot{G_{GSH}} = Q_{GSH} - k_{GSH} * G_{GSH} \; (\boldsymbol{eq. 21})$$

$$Q_r = g_{Q_r} \cdot H^{s-}\left(Q, Q_0, \lambda_{Q,Q_{re}}, n_{Q,Q_{re}}\right) \cdot H^{s+}\left(M, M^0_{Q_{re}}, \lambda_{M,Q_{re}}, n_{M,Q_{re}}\right) \cdot H^{s+}\left(H, H^0_{Q_{re}}, \lambda_{H,Q_{re}}, n_{H,Q_{re}}\right) \cdot H^{s-}\left(A, A^0_{Q_{re}}, \lambda_{A,Q_{re}}, n_{A,Q_{re}}\right) \quad (\boldsymbol{eq.22})$$

$(eqs. 19 - 22)$ **represents the glutamine oxidation rate ($Q_1$), the glutathione synthesis rate ($Q_{GSH}$), the glutathione dynamics ($G_{GSH}$) and the reductive glutamine metabolic rate ($Q_r$) respectively.**

The negative shifted Hill functions $H^{s-}\left(Q, Q_0, \lambda_{Q,Q_1}, n_{Q,Q_1}\right)$, $H^{s-}\left(, Q_0, \lambda_{Q,Q_{GSH}}, n_{Q,Q_{GSH}}\right)$ and $H^{s-}\left(Q, Q_0, \lambda_{Q,Q_{re}}, n_{Q,Q_{re}}\right)$ represent the competition of glutamine oxidation ($Q_1$), glutathione synthesis pathway ($Q_{GSH}$) and reductive glutamine metabolism ($Q_{re}$) on glutamine utilization. The threshold $Q_0$, that is the glutamine uptake rate, in these three shifted Hill functions adds restriction on $Q_1$, $Q_{GSH}$ and $Q_{re}$, since if $Q > Q_0$, that means the glutamine utilization rate is larger than the glutamine uptake rate, these negative shifted Hill functions will decrease $Q_1$, $Q_{GSH}$ and $Q_{re}$ thus decreasing $Q$.

$H^{s+}\left(M, M^0_{Q_1}, \lambda_{M,Q_1}, n_{M,Q_1}\right)$ represents the up-regulation of glutamine oxidation by Myc.
$H^{s-}\left(H, H^0_{Q_1}, \lambda_{H,Q_1}, n_{H,Q_1}\right)$ represents the down-regulation of glutamine oxidation by HIF-1.
$H^{s-}\left(M, M^0_{Q_{GSH}}, \lambda_{M,Q_{GSH}}, n_{M,Q_{GSH}}\right)$ represents the inhibition of glutathione synthesis by Myc.
$H^{s+}\left(M, M^0_{Q_{re}}, \lambda_{M,Q_{re}}, n_{M,Q_{re}}\right)$ represent the up-regulation of reductive glutamine metabolism by Myc.
$\cdot H^{s+}\left(H, H^0_{Q_{re}}, \lambda_{H,Q_{re}}, n_{H,Q_{re}}\right)$ represents the up-regulation of reductive glutamine metabolism by HIF-1.
$H^{s-}\left(A, A^0_{Q_{re}}, \lambda_{A,Q_{re}}, n_{A,Q_{re}}\right)$ represents the down-regulation of reductive glutamine metabolism by AMPK.

$$G_{1,ATP} = 29 * G_1 \quad (\boldsymbol{eq.23})$$

$$G_{2,ATP} = 2 * G_2 \quad (\boldsymbol{eq.24})$$

$$Q_{1,ATP} = 24 * Q_1 \quad (\boldsymbol{eq.25})$$

$$F_{1,ATP} = 106 * F \quad (\boldsymbol{eq.26})$$

$(eqs. 23 - 26)$ **represent the production rates of ATP glucose oxidation ($G_{1,ATP}$) and glycolysis ($G_{2,ATP}$), glutamine oxidation ($Q_{1,ATP}$), FAO ($F_{1,ATP}$), respectively.**

$$F_{2,ATP} = 7 * F_2 \quad (\boldsymbol{eq.27})$$

$$G_{re,ATP} = (15 - 2) * G_{re} \quad (\boldsymbol{eq.28})$$

$$Q_{re,ATP} = 15 * Q_{re} \quad (\boldsymbol{eq.29})$$

$$Q_{GSH,ATP} = 2 * Q_{GSH} \quad (\boldsymbol{eq.30})$$

$(eqs. 27 - 30)$ **represent the ATP consumption rates of reductive fatty acid ($F_{2,ATP}$), reductive glucose ($G_{re,ATP}$) and reductive glutamine ($Q_{re,ATP}$ ) metabolism and GSH synthesis $Q_{GSH,ATP}$), respectively.**

$$X_{ATP} = G_{1,ATP} + G_{2,ATP} + Q_{1,ATP} + F_{1,ATP} - F_{2,ATP} - G_{r,ATP} - Q_{r,ATP} - Q_{GSH,ATP} \quad (\boldsymbol{eq.31})$$

$(eq. 26)$ **represents the net total production rate of ATP ($X_{ATP}$)**.

The definition of the shifted Hill function and the function $C_{R_{nox}}^{comp}$ representing the competitive regulation of noxROS by AMPK and HIF-1 are as follows.

The shifted Hill function (1) $H^s(X, X_0, \lambda, n)$ is defined top be

$H^s(X, X_0, \lambda, n) = \frac{1+\lambda\left(\frac{X}{X_0}\right)^n}{1+\left(\frac{X}{X_0}\right)^n}$, where $X$ represents the level of the regulator, $X_0$ represents the threshold, $\lambda$ represents the fold-change and $n$ represents the Hill coefficient.

$H^{s+}(X, X_0, \lambda, n) = \frac{1+\lambda^+\left(\frac{X}{X_0}\right)^n}{1+\left(\frac{X}{X_0}\right)^n}$, where $\lambda^+ > 1$, representing the excitatory regulation.

$H^{s-}(X, X_0, \lambda, n) = \frac{1+\lambda^-\left(\frac{X}{X_0}\right)^n}{1+\left(\frac{X}{X_0}\right)^n}$, where $\lambda^- < 1$, representing the inhibitory regulation.

The competitive regulations of noxROS by AMPK and HIF-1 ($C^{comp}$) (2) is defined as follows:

$C_{R_{nox}}^{comp}\left(g_0, H, g_{H,R_{nox}}, H_{R_{nox}}^0, n_{H,R_{nox}}, A, g_{A,R_{nox}}, A_{R_{nox}}^0, n_{A,R_{nox}}\right) =$

$\frac{g_0+g_{H,R_{nox}}\left(\frac{H}{H_{R_{nox}}^0}\right)^{n_{H,R_{nox}}}+g_{A,R_{nox}}\left(\frac{A}{A_{R_{nox}}^0}\right)^{n_{A,R_{nox}}}}{1+\left(\frac{H}{H_{R_{nox}}^0}\right)^{n_{H,R_{nox}}}+\left(\frac{A}{A_{R_{nox}}^0}\right)^{n_{A,R_{nox}}}}$, where $g_0 = 1$ representing the basal noxROS production, $g_{H,R_{nox}}\left(\frac{H}{H_{R_{nox}}^0}\right)^{n_{H,R_{nox}}}$ represents the regulation of noxROS production by HIF-1 and $g_{A,R_{nox}}\left(\frac{A}{A_{R_{nox}}^0}\right)^{n_{A,R_{nox}}}$ represents the regulation of noxROS production by AMPK.

The baseline model parameters are presented in **Table 1**

**Data Analysis**

PanCancer Atlas data, including the clinical data were downloaded from The cBio Cancer Genomics Portal (cBioPortal, https://www.cbioportal.org/ ), the analyzed cancers were: hepatocellular carcinoma, breast invasive carcinoma, kidney renal clear cell carcinoma, lung adenocarcinoma, lung squamous cell carcinoma, colorectal adenocarcinoma, skin cutaneous melanoma, acute myeloid leukemia, and prostate adenocarcinoma.

Data analysis and visualization were performed using several R packages. Heat maps and clustering analyses were conducted using the `ComplexHeatmap` package (Elis R, 2016). The `ggplot2` package was utilized for data visualization, and the `ggpubr` package was employed for statistical analysis, including the t-test. For the Kaplan-Meier plots, patient data were categorized based on the results of the clustering analysis. The overall survival analysis was conducted using the log-rank test. The `survival` package (Therneau and Grambsch, 2000) was used for survival analysis, and the `survminer` package (Kassambara, Kosinski, and Biecek, 2021) was used to create the Kaplan-Meier plots.

### Table 1. The values of parameters

| Parameters | Value | Unit | Description |
|---|---|---|---|
| $g_{R_{mt}}$ | 50 | $nmol \cdot L^{-1} \cdot h^{-1}$ | Production rate of mtROS |
| $\gamma_{G_1}$ | 1 | - | mtROS production in glucose oxidation |
| $\gamma_F$ | 9/2 | - | mtROS production in fatty acid oxidation |
| $\gamma_{Q_1}$ | 5 | - | mtROS production in glutamine oxidation |
| $k_{R_{mt}}$ | 5 | $min^{-1}$ | Degradation rate of mtROS |
| $A^0_{R_{mt}}$ | 350 | $nmol \cdot L^{-1}$ | Threshold of mtROS inhibition by AMPK |
| $\lambda_{A,R_{mt}}$ | 2 | - | Fold change of mtROS inhibition by AMPK |
| $n_{A,R_{mt}}$ | 2 | - | Hill coefficient |
| $G^0_{GSH,R}$ | 10 | $\mu mol \cdot L^{-1} \cdot s^{-1}$ | Threshold of mtROS clearance by AMPK |
| $\lambda_{GSH,R}$ | 1.2 | - | Fold change of mtROS clearance by AMPK |
| $n_{GSH,R}$ | 2 | - | Hill coefficient |
| $g_{R_{nox}}$ | 40 | $\mu mol \cdot L^{-1} \cdot min^{-1}$ | Production rate of noxROS |
| $g_0$ | 1 | - | Basal noxROS |
| $g_{H,R_{nox}}$ | 5 | - | Fold-change of noxROS activation by HIF-1 |
| $H^0_{R_{nox}}$ | 250 | $nmol \cdot L^{-1}$ | Threshold of noxROS activation by HIF-1 |
| $n_{H,R_{nox}}$ | 2 | - | Hill coefficient |
| $g_{A,R_{nox}}$ | 0.2 | - | Fold change of noxROS inhibition by AMPK |
| $A^0_{R_{nox}}$ | 150 | $nmol \cdot L^{-1}$ | Threshold of noxROS inhibition by AMPK |
| $n_{A,R_{nox}}$ | 2 | - | Hill coefficient |
| $k_{R_{nox}}$ | 5 | $min^{-1}$ | Degradation rate of noxROS |
| $g_A$ | 40 | $nmol \cdot L^{-1} \cdot h^{-1}$ | Production rate of AMPK |
| $R^0_{T,A}$ | 250 | $\mu mol \cdot L^{-1}$ | Threshold of AMPK activation by ROS |
| $\lambda_{R_T,A}$ | 8 | - | Fold change of AMPK activation by ROS |

| | | | |
|---|---|---|---|
| $n_{R_T,A}$ | 4 | - | Hill coefficient |
| $H_A^0$ | 150 | $nmol \cdot L^{-1}$ | Threshold of AMPK inhibition by HIF-1 |
| $\lambda_{H,A}$ | 0.1 | - | Fold-change of AMPK inhibition by HIF-1 |
| $n_{H,A}$ | 1 | - | Hill coefficient |
| $\lambda_{X_{ATP},A}$ | 0.25 | - | Fold-change of AMPK inhibition by ATP |
| $X_{ATP,A}^0$ | 2000 | $\mu mol \cdot L^{-1} \cdot s^{-1}$ | Threshold of AMPK inhibition by ATP |
| $n_{X_{ATP},A}$ | 2 | - | Hill coefficient |
| $k_A$ | 0.2 | $h^{-1}$ | Degradation rate of AMPK |
| $g_H$ | 15 | $nmol \cdot L^{-1} \cdot h^{-1}$ | Production rate of HIF-1 |
| $A_H^0$ | 150 | $nmol \cdot L^{-1}$ | Threshold of HIF-1 inhibition by AMPK |
| $\lambda_{A,H}$ | 0.1 | - | Fold-change of HIF-1 inhibition by AMPK |
| $n_{A,H}$ | 1 | - | Hill coefficient |
| $k_H$ | 0.25 | $h^{-1}$ | Degradation rate of HIF-1 |
| $G_{2,H}^0$ | 250 | $\mu mol \cdot L^{-1} \cdot s^{-1}$ | Threshold of HIF-1 activation by glycolysis |
| $\lambda_{G_2,H}$ | 0.1 | - | Fold-change of HIF-1 activation by glycolysis |
| $n_{G_2,H}$ | 4 | - | Hill coefficient |
| $M_{M,H}^0$ | 150 | $\mu mol \cdot L^{-1} \cdot s^{-1}$ | Threshold of MYC inhibting HIF-1 degradation |
| $\lambda_{M,H}$ | 0.8 | - | Fold change of MYC inhibting HIF-1 degradation |
| $n_{M,H}$ | 2 | - | Hill coefficient |
| $\lambda_{R_T,H}$ | 0.2 | - | Fold-change of HIF-1 stabilization by ROS |
| $R_{T,H}^0$ | 40 | $\mu mol \cdot L^{-1}$ | Threshold of HIF-1 stabilization by ROS |
| $n_{R_T,H}$ | 4 | - | Hill coefficient |
| $g_{H,G_0}$ | 20 | $\mu mol \cdot L^{-1} \cdot s^{-1}$ | Basal glucose uptake rate |
| $H_{G_0}^0$ | 150 | $nmol \cdot L^{-1}$ | Threshold of glucose uptake regulated by HIF-1 |
| $\lambda_{H,G_0}$ | 6 | - | Fold-change of glucose uptake regulated by HIF-1 |

| | | | |
|---|---|---|---|
| $n_{H,G_0}$ | 4 | - | Hill coefficient |
| $g_{A,G_0}$ | 20 | $\mu mol \cdot L^{-1} \cdot s^{-1}$ | Basal glucose uptake rate |
| $A_{G_0}^0$ | 200 | $nmol \cdot L^{-1}$ | Threshold of glucose uptake regulated by AMPK |
| $\lambda_{A,G_0}$ | 4 | - | Fold-change of glucose uptake regulated by AMPK |
| $n_{A,G_0}$ | 2 | - | Hill coefficient |
| $M_{G_0}^0$ | 150 | $\mu mol \cdot L^{-1} \cdot s^{-1}$ | Threshold of glucose uptake regulated by MYC |
| $\lambda_{M,G_0}$ | 2 | - | Fold change of glucose uptake regulated by MYC |
| $n_{M,G_0}$ | 2 | - | Hill coefficient |
| $M_{Q_0}^0$ | 150 | $\mu mol \cdot L^{-1} \cdot s^{-1}$ | Threshold of glutamine uptake regulated by MYC |
| $\lambda_{M,Q_0}$ | 2 | - | Fold-change of glutamine uptake regulated by MYC |
| $n_{M,Q_0}$ | 2 | - | Fold-change of glucose uptake regulated by MYC |
| $g_{A,C_0}$ | 15 | $\mu mol \cdot L^{-1} \cdot s^{-1}$ | Basal utilization rate of acetyl-CoA for OXPHOS |
| $A_{C_0}^0$ | 250 | $nmol \cdot L^{-1}$ | Threshold of acetyl-CoA utilization regulated by AMPK |
| $\lambda_{A,C_0}$ | 8 | - | Fold-change of acetyl-CoA utilization regulated by AMPK |
| $n_{A,C_0}$ | 4 | - | Hill coefficient |
| $g_{G_1}$ | 100 | $\mu mol \cdot L^{-1} \cdot s^{-1}$ | Basal glucose oxidation rate |
| $\lambda_{G,G_1}$ | 0.1 | - | Restriction of glucose oxidation rate by glucose uptake rate |
| $n_{G,G_1}$ | 2 | - | Hill coefficient |
| $\lambda_{C,G_1}$ | 0.1 | - | Restriction of glucose oxidation by acetyl-CoA utilization rate |
| $n_{C,G_1}$ | 4 | - | Hill coefficient |
| $g_{G_2}$ | 150 | $\mu mol \cdot L^{-1} \cdot s^{-1}$ | Basal glycolysis rate |
| $\lambda_{G,G_2}$ | 0.1 | - | Restriction of glycolysis rate by glucose uptake rate |
| $n_{G,G_2}$ | 2 | - | Hill coefficient |
| $H_{G_2}^0$ | 200 | $nmol \cdot L^{-1}$ | Threshold of glycolysis upregulation by HIF-1 |
| $\lambda_{H,G_2}$ | 8 | - | Fold-change of glycolysis upregulation by HIF-1 |

| | | | |
|---|---|---|---|
| $n_{H,G_2}$ | 4 | - | Hill coefficient |
| $g_{G_r}$ | 60 | $nmol \cdot L^{-1} \cdot h^{-1}$ | Basal reductive glucose metabolic rate |
| $\lambda_{G,G_{re}}$ | 0.1 | - | Restriction of reductive glucose metabolic rate by glucose uptake rate |
| $n_{G,G_{re}}$ | 2 | - | Hill coefficient |
| $A^0_{G_{re}}$ | 200 | $\mu mol \cdot L^{-1} \cdot s^{-1}$ | Threshold of reductive glucose metabolism regulation by AMPK |
| $\lambda_{A,G_{re}}$ | 0.25 | - | Fold-change of reductive glucose metabolism regulation by AMPK |
| $n_{A,G_{re}}$ | 2 | - | Hill coefficient |
| $M^0_{G_{re}}$ | 150 | $\mu mol \cdot L^{-1} \cdot s^{-1}$ | Threshold of reductive glucose metabolism regulation by MYC |
| $\lambda_{M,G_{re}},$ | 2 | - | Fold-change of reductive glucose metabolism regulation by MYC |
| $n_{M,G_{re}}$ | 2 | - | Hill coefficient |
| $g_f$ | 2 | $\mu mol \cdot L^{-1} \cdot s^{-1}$ | Basal FAO rate |
| $\lambda_{C,F}$ | 0.1 | - | Restriction of FAO rate by acetyl-CoA utilization rate |
| $n_{C,F}$ | 2 | - | Hill coefficient |
| $A^0_{F_1}$ | 200 | $nmol \cdot L^{-1}$ | Threshold of FAO upregulation by AMPK |
| $\lambda_{A,F_1}$ | 6 | - | Fold-change of FAO upregulation by AMPK |
| $n_{A,F_1}$ | 4 | - | Hill coefficient |
| $g_{F0}$ | 5 | $mol \cdot L^{-1} \cdot s^{-1}$ | Basic fatty acid update rate |
| $A^0_{F_0}$ | 200 | $\mu mol \cdot L^{-1} \cdot s^{-1}$ | Threshold of fatty acid uptake regulation by AMPK |
| $n_{A,F_0}$ | 2 | - | Hill coefficient |
| $\lambda_{A,F_0}$ | 4 | - | Fold-change of fatty acid uptake regulation by AMPK |
| $\lambda_{F,F_1}$ | 0.8 | - | Restriction of FAO rate by fatty acid uptake rate |
| $n_{F,F_1}$ | 2 | - | Hill coefficient |
| $H^0_F$ | 200 | $\mu mol \cdot L^{-1} \cdot s^{-1}$ | Threshold of FAO regulation by HIF-1 |
| $\lambda_{H,F},$ | 0.8 | - | Fold-change of FAO regulation by HIF-1 |
| $n_{H,F}$ | 4 | - | Hill coefficient |

| | | | |
|---|---|---|---|
| $\gamma_{G,F}$ | 0.1 | - | Lipogenesis rate of glucose |
| $\gamma_{G,Q}$ | 0.1 | - | Lipogenesis rate of glutamine |
| $g_{F_r}$ | 75 | $nmol \cdot L^{-1} \cdot h^{-1}$ | Basal reductive fatty acid metabolic rate |
| $\lambda_{F,F_r}$ | 0.1 | - | Restriction of FAO rate by fatty acid uptake rate |
| $n_{F,F_r}$ | 2 | - | Hill coefficient |
| $g_{Q_1}$ | 30 | $nmol \cdot L^{-1} \cdot h^{-1}$ | Basal glutamine oxidation rate |
| $H_{Q_1}^0$ | 350 | $\mu mol \cdot L^{-1} \cdot s^{-1}$ | Threshold of glutamine oxidation regulation by HIF-1 |
| $\lambda_{H,Q_1}$ | 0.2 | - | Fold change of glutamine oxidation regulation by HIF-1 |
| $n_{H,Q_1}$ | 4 | - | Hill coefficient |
| $\lambda_{Q,Q_1}$ | 0.5 | - | Restriction of glutamine oxidation rate by glutamine uptake rate |
| $n_{Q,Q_1}$ | 2 | - | Hill coefficient |
| $g_{Q_0}$ | 10 | $nmol \cdot L^{-1} \cdot h^{-1}$ | Basal glutamine update rate |
| $M_{Q_0}^0$ | 150 | $nmol \cdot L^{-1}$ | Threshold of glutamine uptake regulation by MYC |
| $\lambda_{M,Q_0}$ | 2 | - | Fold change of glutamine uptake regulation by MYC |
| $n_{M,Q_0}$ | 2 | - | Hill coefficient |
| $M_{Q_1}^0$ | 150 | $\mu mol \cdot L^{-1} \cdot s^{-1}$ | Threshold of glutamine oxidation regulation by MYC |
| $\lambda_{M,Q_1}$ | 2 | - | Fold change of glutamine oxidation regulation by MYC |
| $n_{M,Q_1}$ | 2 | - | Hill coefficient |
| $g_{Q_{GSH}}$ | 30 | $nmol \cdot L^{-1} \cdot h^{-1}$ | Basal glutathione synthesis rate |
| $\lambda_{Q,Q_{GSH}}$ | 0.5 | - | Restriction of glutathione synthesis rate by glutamine uptake rate |
| $n_{Q,Q_{GSH}}$ | 2 | - | Hill coefficient |
| $M_{Q_{GSH}}^0$ | 150 | $\mu mol \cdot L^{-1} \cdot s^{-1}$ | Threshold of glutathione synthesis regulation by MYC |
| $\lambda_{M,Q_{GSH}}$ | 0.9 | - | Fold change of glutathione synthesis regulation by MYC |
| $n_{M,Q_{GSH}}$ | 2 | - | Hill coefficient |
| $k_{GSH}$ | 1 | $h^{-1}$ | Decay rate of glutathione |

| | | | |
|---|---|---|---|
| $g_{Q_r}$ | 20 | $nmol \cdot L^{-1} \cdot h^{-1}$ | Basal reductive glutamine metabolic rate |
| $\lambda_{Q,Q_r}$ | 0.5 | - | Restriction of reductive glutamine metabolic rate by glutamine uptake rate |
| $n_{Q,Q_r}$ | 2 | - | Hill coefficient |
| $M_{Q_r}^0$ | 150 | $\mu mol \cdot L^{-1} \cdot s^{-1}$ | Threshold of reductive glutamine metabolism regulation by MYC |
| $\lambda_{M,Q_r}$ | 2 | - | Fold change of reductive glutamine metabolism regulation by MYC |
| $n_{M,Q_r}$ | 2 | - | Hill coefficient |
| $H_{Q_r}^0$ | 200 | $\mu mol \cdot L^{-1} \cdot s^{-1}$ | Threshold of reductive glutamine metabolism regulation by HIF-1 |
| $\lambda_{H,Q_r}$ | 2 | - | Fold change of reductive glutamine metabolism regulation by HIF-1 |
| $n_{H,Q_r}$ | 4 | - | Hill coefficient |
| $A_{Q_r}^0$ | 200 | $\mu mol \cdot L^{-1} \cdot s^{-1}$ | Threshold of reductive glutamine metabolism regulation by AMPK |
| $\lambda_{A,Q_r}$ | 0.5 | - | Fold change of reductive glutamine metabolism regulation by AMPK |
| $n_{A,Q_r}$ | 2 | - | Hill coefficient |

**Supplementary Information**

**Supplementary Figure 1**

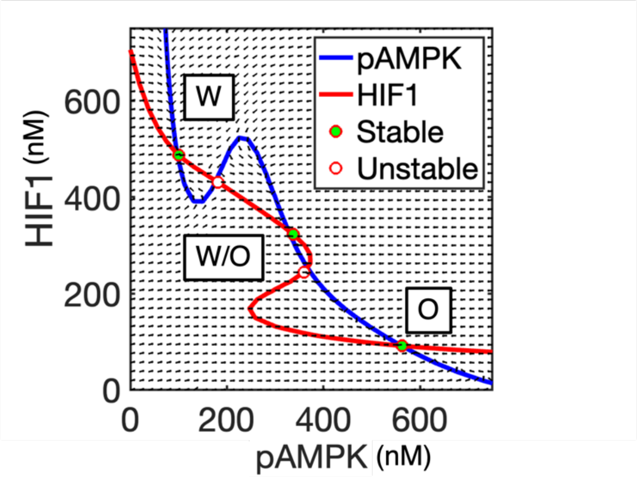


**Supplementary Figure 1: Nullclines and steady states in the phase space of AMPK and HIF-1 when the cells have high MYC levels.** Nullclines and steady states in the phase space of AMPK and HIF-1. The red line represents the nullcline where the rate of change of HIF-1 (*dH/dt*) is zero, and the blue line represents the nullcline where the rate of change of AMPK (*dA/dt*) is zero. Solid dots represent stable steady states while hollow dots represent unstable steady states. Each stable state is associated with a metabolic phenotype. With a low MYC expression, cells can acquire the Warburg phenotype, referred to as the state “W”, an OXPHOS phenotype, referred to as the state “O”, and a hybrid phenotype with intermediate activities of AMPK and HIF-1 called the state “W/O”.

**Supplementary Figure 2**

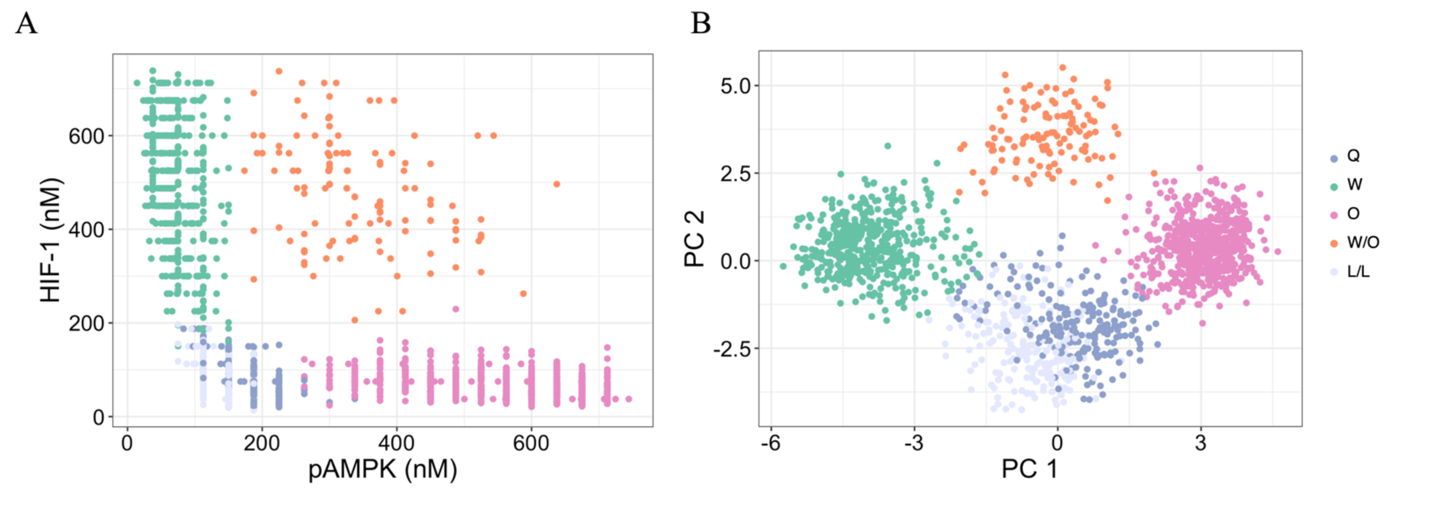


**Supplementary Figure 2. A.** Plot of pAMPK vs HIF-1 illustrating the relationship between both components' levels across all generated steady states. **B**. A scatter plot showing the results of a Principal Component Analysis (PCA). Each point in the plot represents a steady state, plotted according to its scores on the first two principal components (PC1 and PC2). Different clusters are indicated by different colors, providing a visual representation of the grouping of steady states in the reduced-dimensional space of the PCA. Both plots, A and B, show the state annotations for the 'L/L' cluster, indicating the position of the Low Metabolic State. These are considered low due to their glucose oxidation, glycolysis, and Fatty acid Oxidation values having Z-scores less than 0, respectively.

**Supplementary Figure 3**

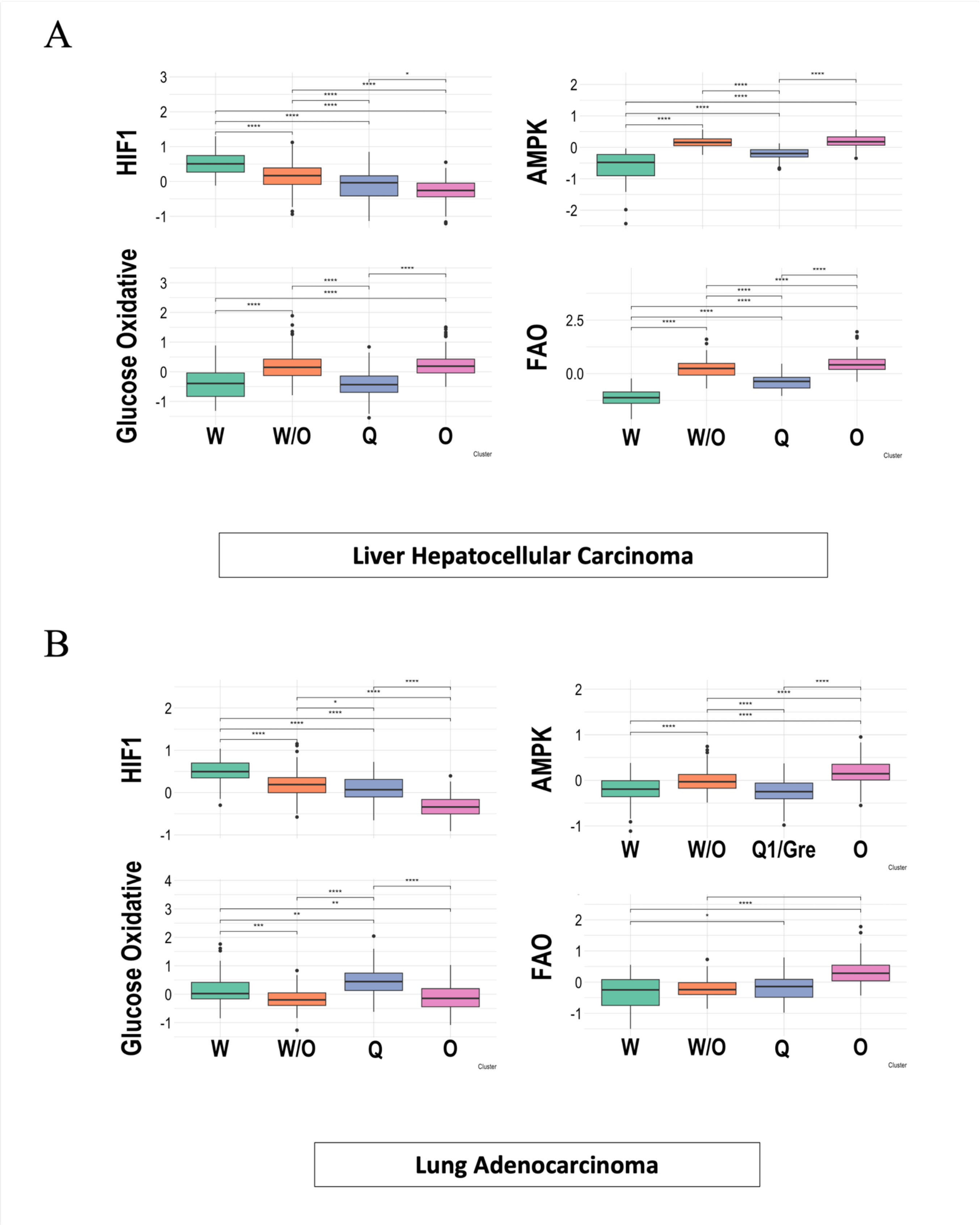


**Supplementary Figure 3.** Box plots summarizing and showing the differences of the scores according to each identified cluster for Liver hepatocellular carcinoma (**A**) and lung adenocarcinoma (**B**). A t-test was used to test the significance of each pair of clusters.

**Supplementary Figure 4**

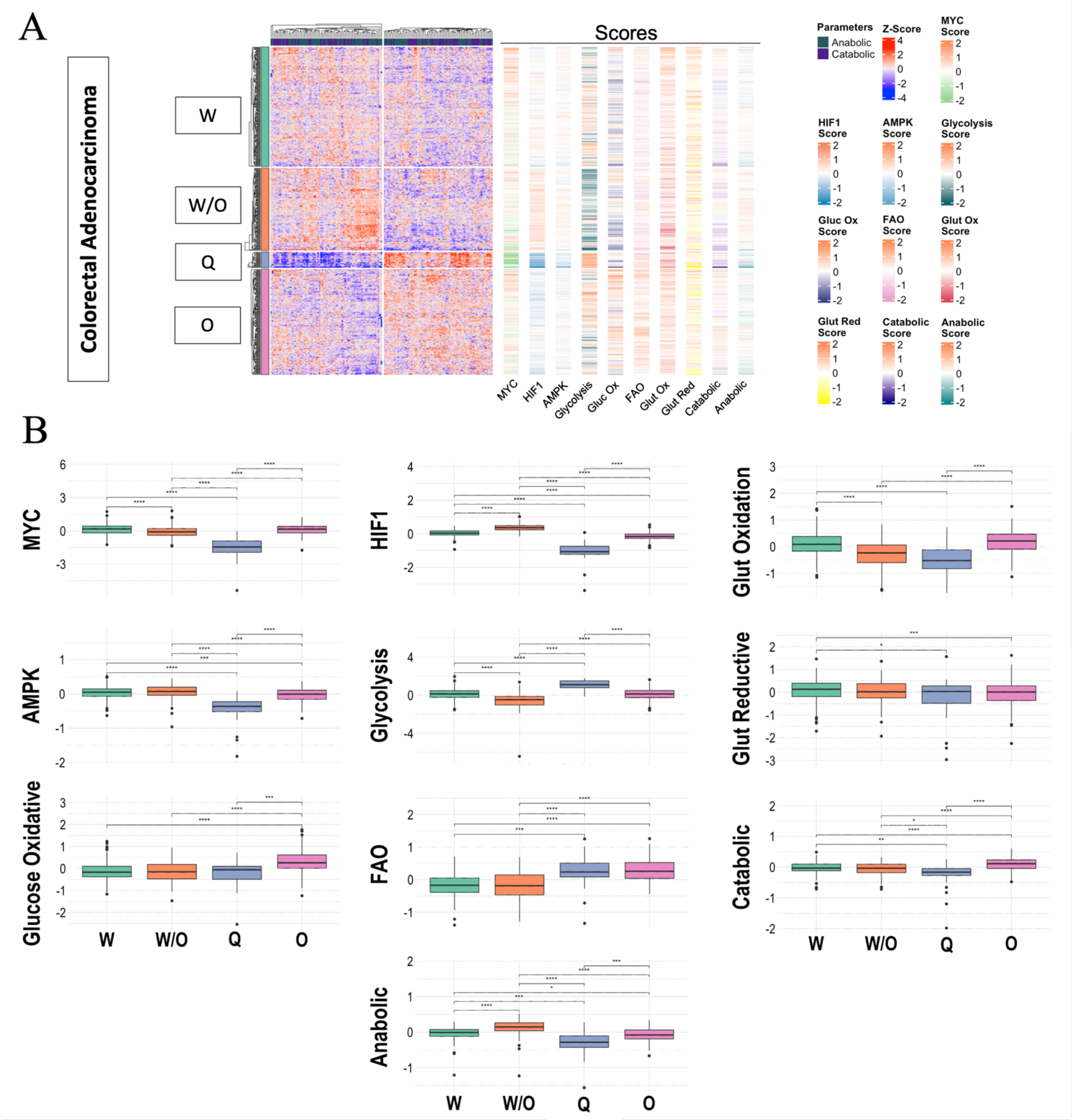


**Supplementary Figure 4. The association between gene activity and metabolic pathway activity for colorectal carcinoma**. **A.** Heatmap of RNA-seq data. Each row represents a patient sample, and each column represents the expression of selected genes, which are divided according to their metabolic or anabolic activities. Four distinct clusters can be identified in the heatmap. The adjacent one-column heatmaps represent the scores for MYC, HIF-1, AMPK, Glycolysis, Glucose Oxidative, FAO, Glutamine Oxidative, Glutamine Reductive, Catabolic, and Anabolic activities, respectively. **B.** Box plots summarizing and showing the differences of the scores according to each identified cluster. A t-test was used to test the significance of each pair of clusters.

**Supplementary Figure 5**

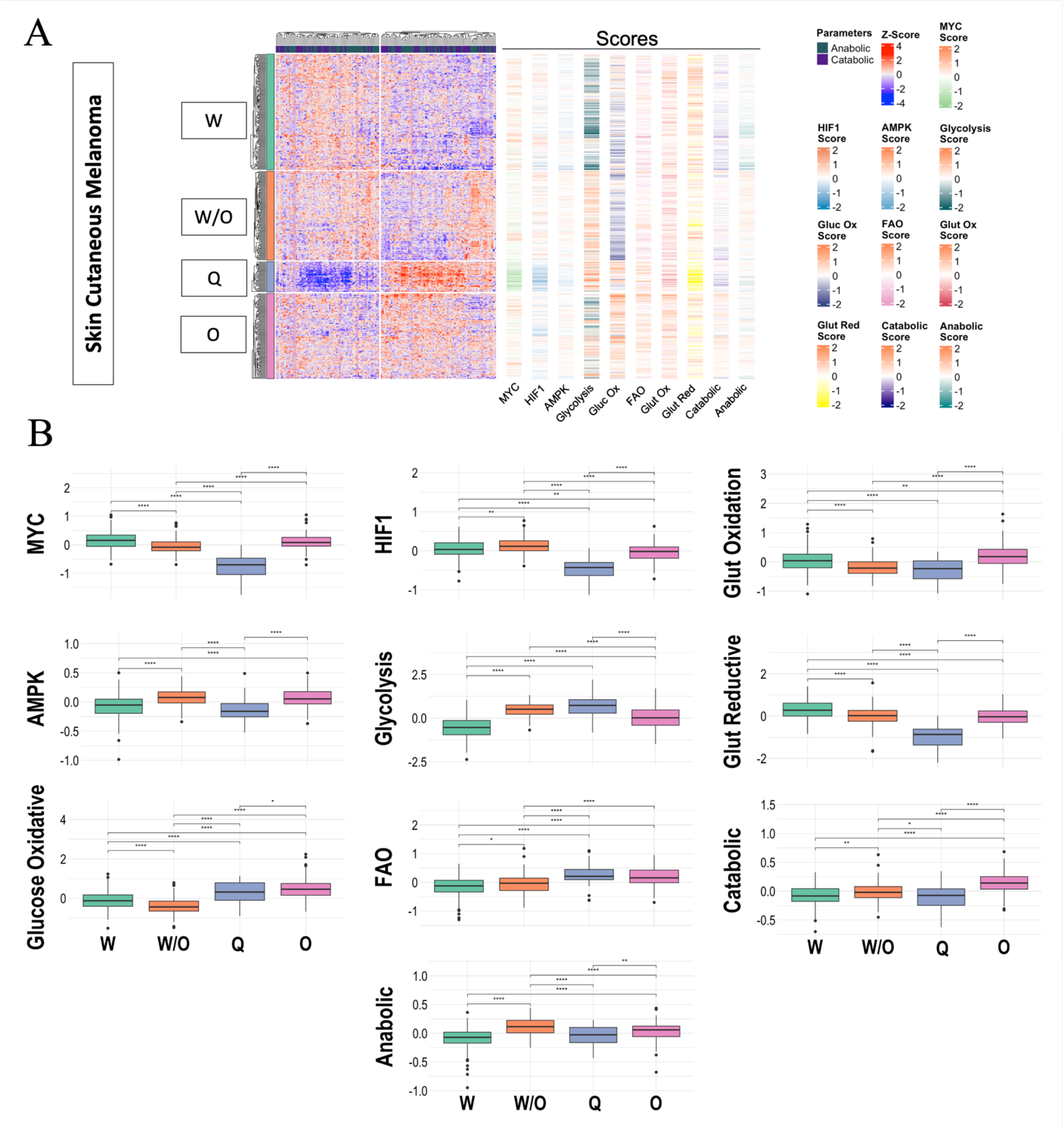


**Supplementary Figure 5. The association between gene activity and metabolic pathway activity for skin cutaneous melanoma**. **A.** Heatmap of RNA-seq data. Each row represents a patient sample, and each column represents the expression of selected genes, which are divided according to their metabolic or anabolic activities. Four distinct clusters can be identified in the heatmap. The adjacent one-column heatmaps represent the scores for MYC, HIF-1, AMPK, Glycolysis, Glucose Oxidative, FAO, Glutamine Oxidative, Glutamine Reductive, Catabolic, and Anabolic activities, respectively. **B.** Box plots summarizing and showing the differences of the scores according to each identified cluster. A t-test was used to test the significance of each pair of clusters.

**Supplementary Figure 6**

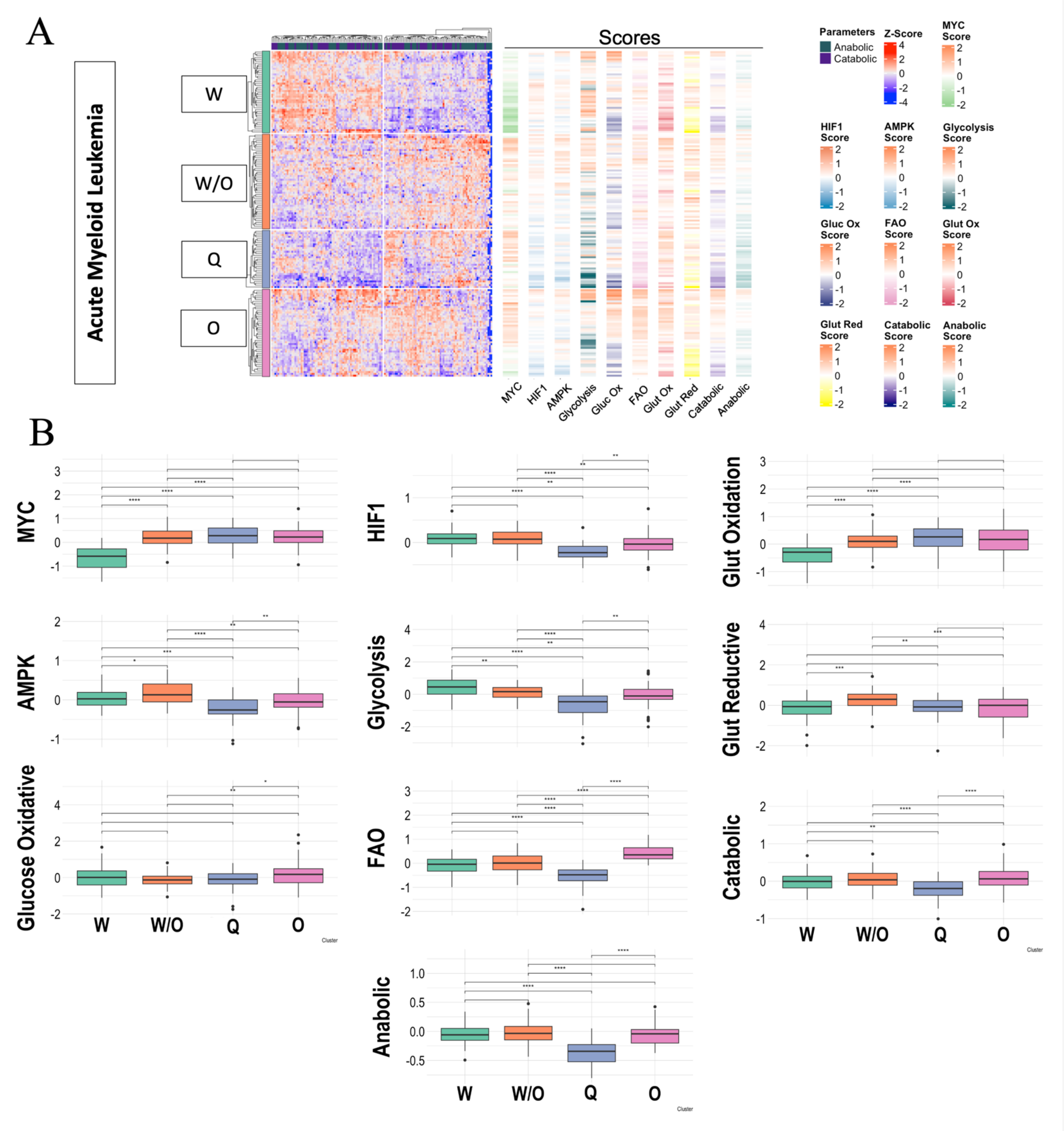


**Supplementary Figure 6. The association between gene activity and metabolic pathway activity for acute myeloid leukemia**. **A.** Heatmap of RNA-seq data. Each row represents a patient sample, and each column represents the expression of selected genes, which are divided according to their metabolic or anabolic activities. Four distinct clusters can be identified in the heatmap. The adjacent one-column heatmaps represent the scores for MYC, HIF-1, AMPK, Glycolysis, Glucose Oxidative, FAO, Glutamine Oxidative, Glutamine Reductive, Catabolic, and Anabolic activities, respectively. **B.** Box plots summarizing and showing the differences of the scores according to each identified cluster. A t-test was used to test the significance of each pair of clusters.

**Supplementary Figure 7**

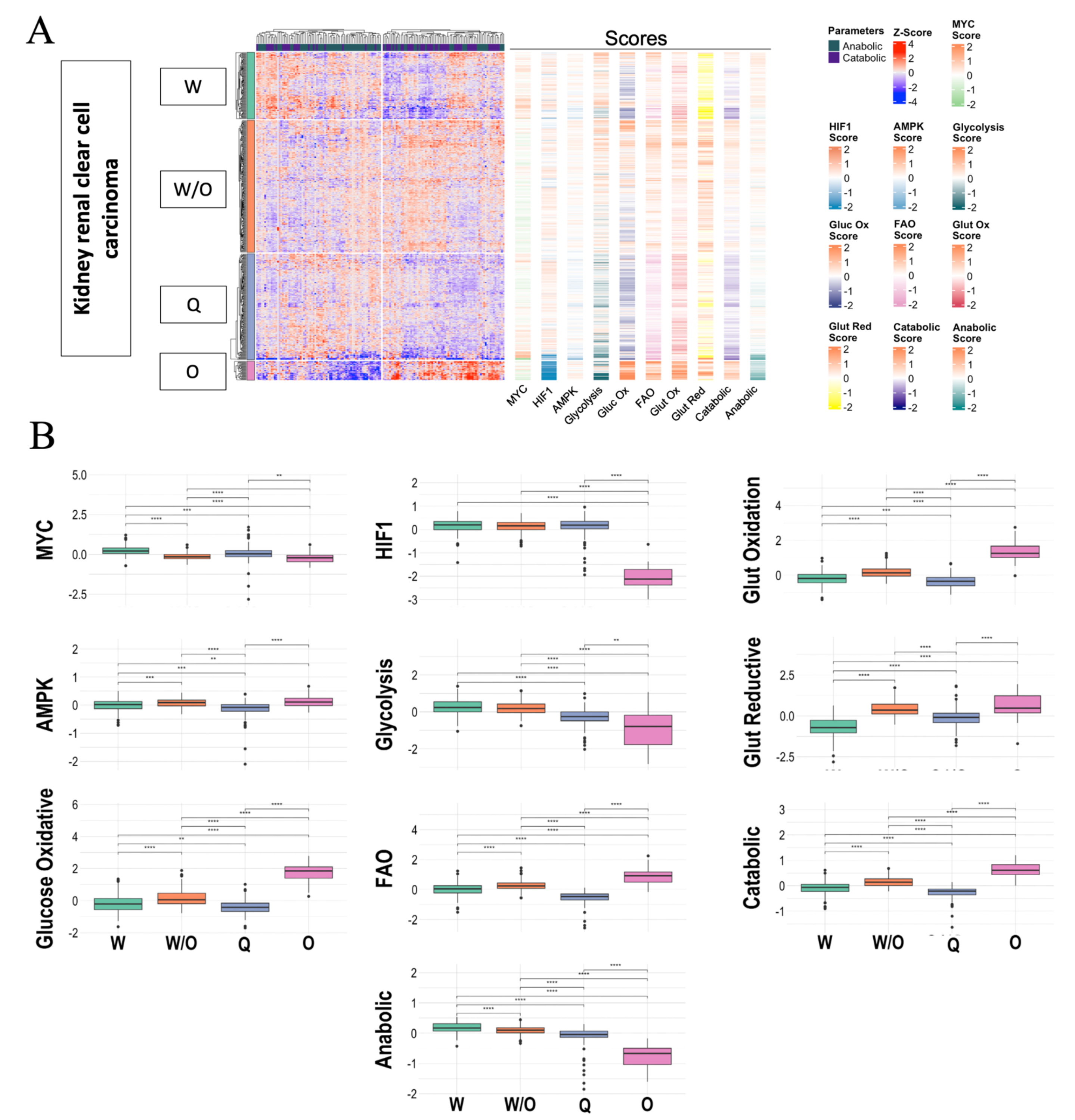


**Supplementary Figure 7. The association between gene activity and metabolic pathway activity for Kidney renal clear cell carcinoma**. **A.** Heatmap of RNA-seq data. Each row represents a patient sample, and each column represents the expression of selected genes, which are divided according to their metabolic or anabolic activities. Four distinct clusters can be identified in the heatmap. The adjacent one-column heatmaps represent the scores for MYC, HIF-1, AMPK, Glycolysis, Glucose Oxidative, FAO, Glutamine Oxidative, Glutamine Reductive, Catabolic, and Anabolic activities, respectively. **B.** Box plots summarizing and showing the differences of the scores according to each identified cluster. A t-test was used to test the significance of each pair of clusters.

**Supplementary Figure 8**

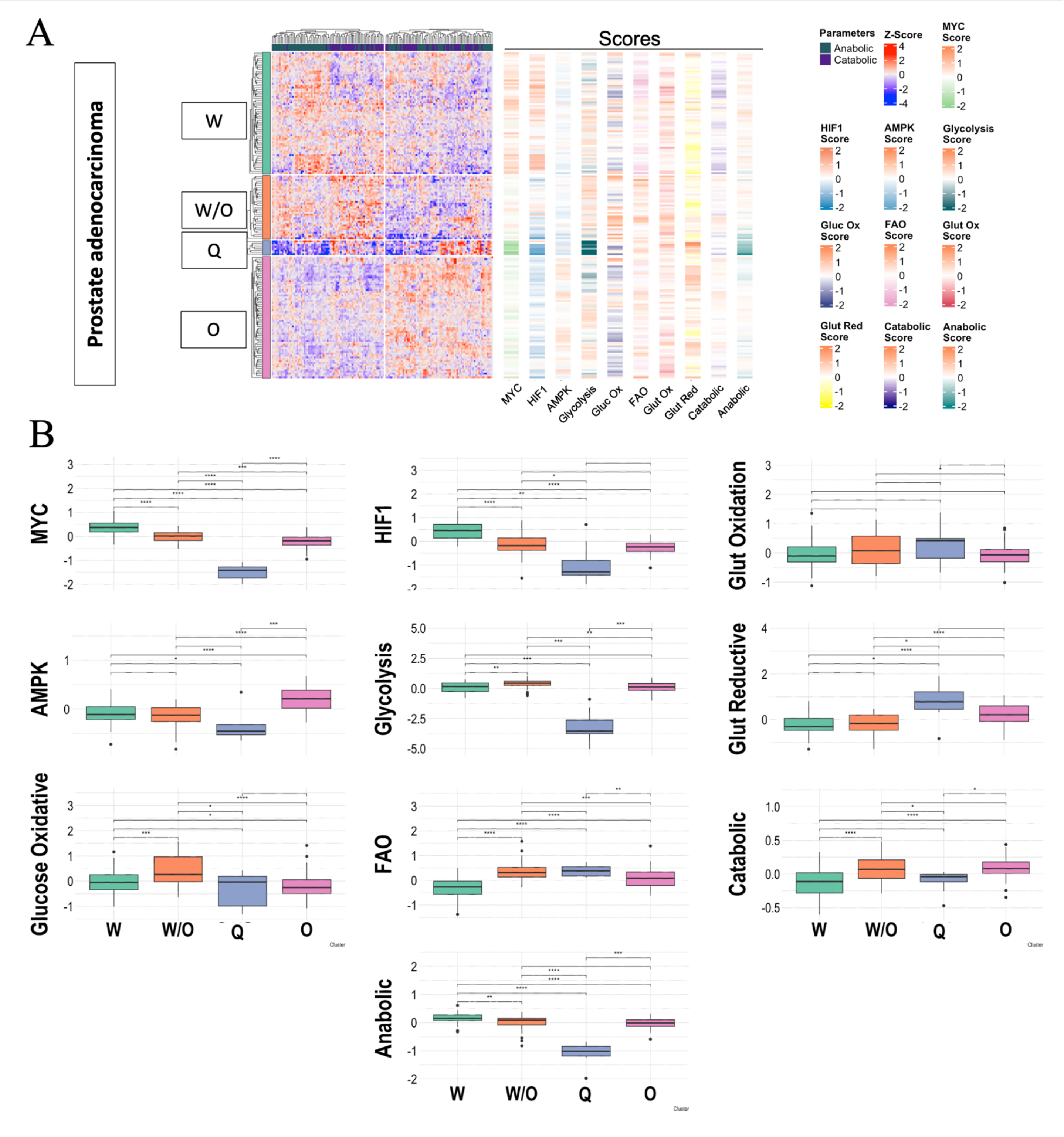


**Supplementary Figure 8. The association between gene activity and metabolic pathway activity for prostate adenocarcinoma**. **A.** Heatmap of RNA-seq data. Each row represents a patient sample, and each column represents the expression of selected genes, which are divided according to their metabolic or anabolic activities. Four distinct clusters can be identified in the heatmap. The adjacent one-column heatmaps represent the scores for MYC, HIF-1, AMPK, Glycolysis, Glucose Oxidative, FAO, Glutamine Oxidative, Glutamine Reductive, Catabolic, and Anabolic activities, respectively. **B.** Box plots summarizing and showing the differences of the scores according to each identified cluster. A t-test was used to test the significance of each pair of clusters.

**Supplementary Figure 9**

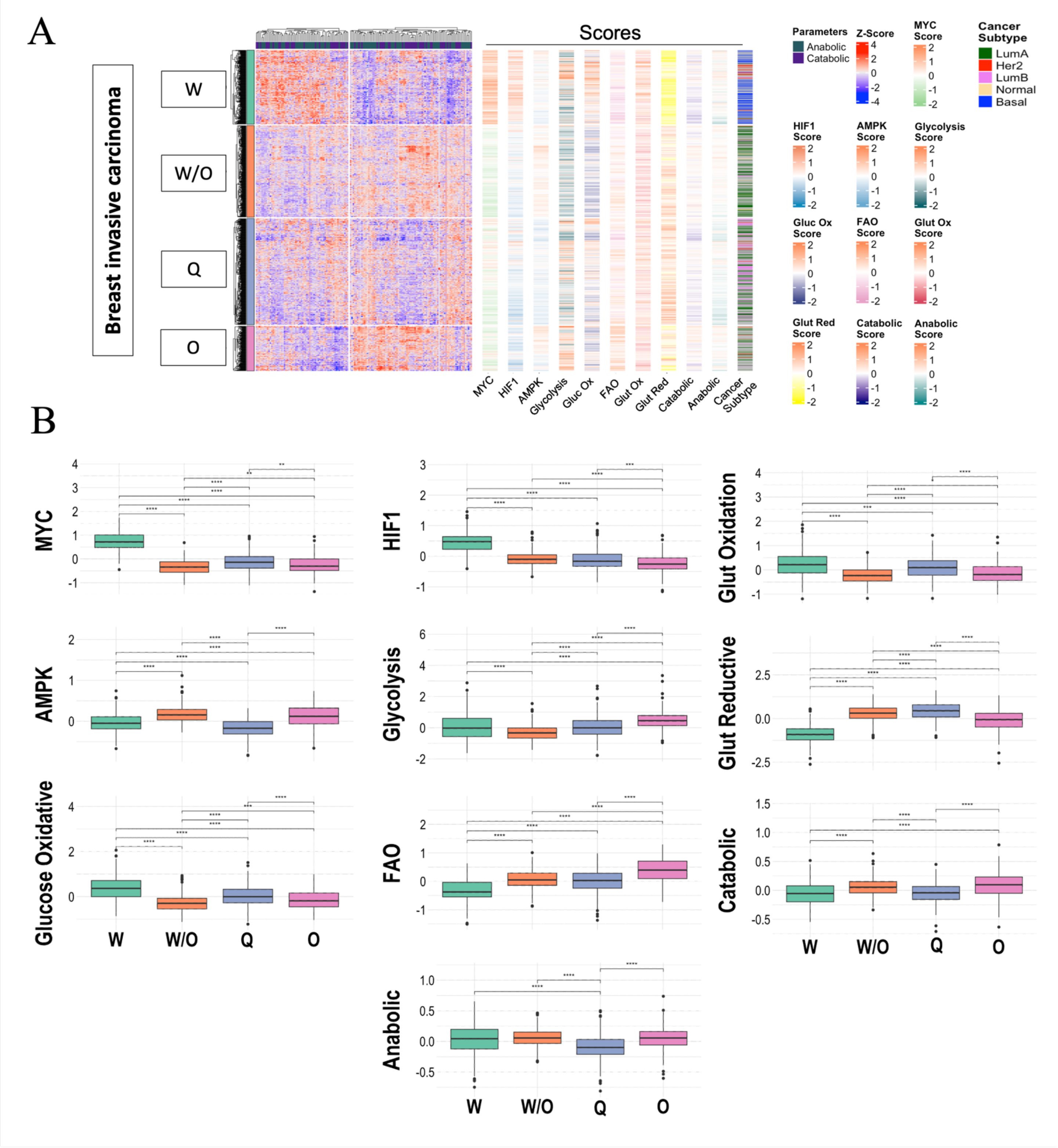


**Supplementary Fig 9: The association between gene activity and metabolic pathway activity for breast invasive carcinoma**. **A.** Heatmap of RNA-seq data. Each row represents a patient sample, and each column represents the expression of selected genes, which are divided according to their metabolic or anabolic activities. Four distinct clusters can be identified in the heatmap. The adjacent one-column heatmaps represent the scores for MYC, HIF-1, AMPK, Glycolysis, Glucose Oxidative, FAO, Glutamine Oxidative, Glutamine Reductive, Catabolic, and Anabolic activities, respectively. **B.** Box plots summarizing and showing the differences of the scores according to each identified cluster. A t-test was used to test the significance of each pair of clusters.

**Supplementary Figure 10**

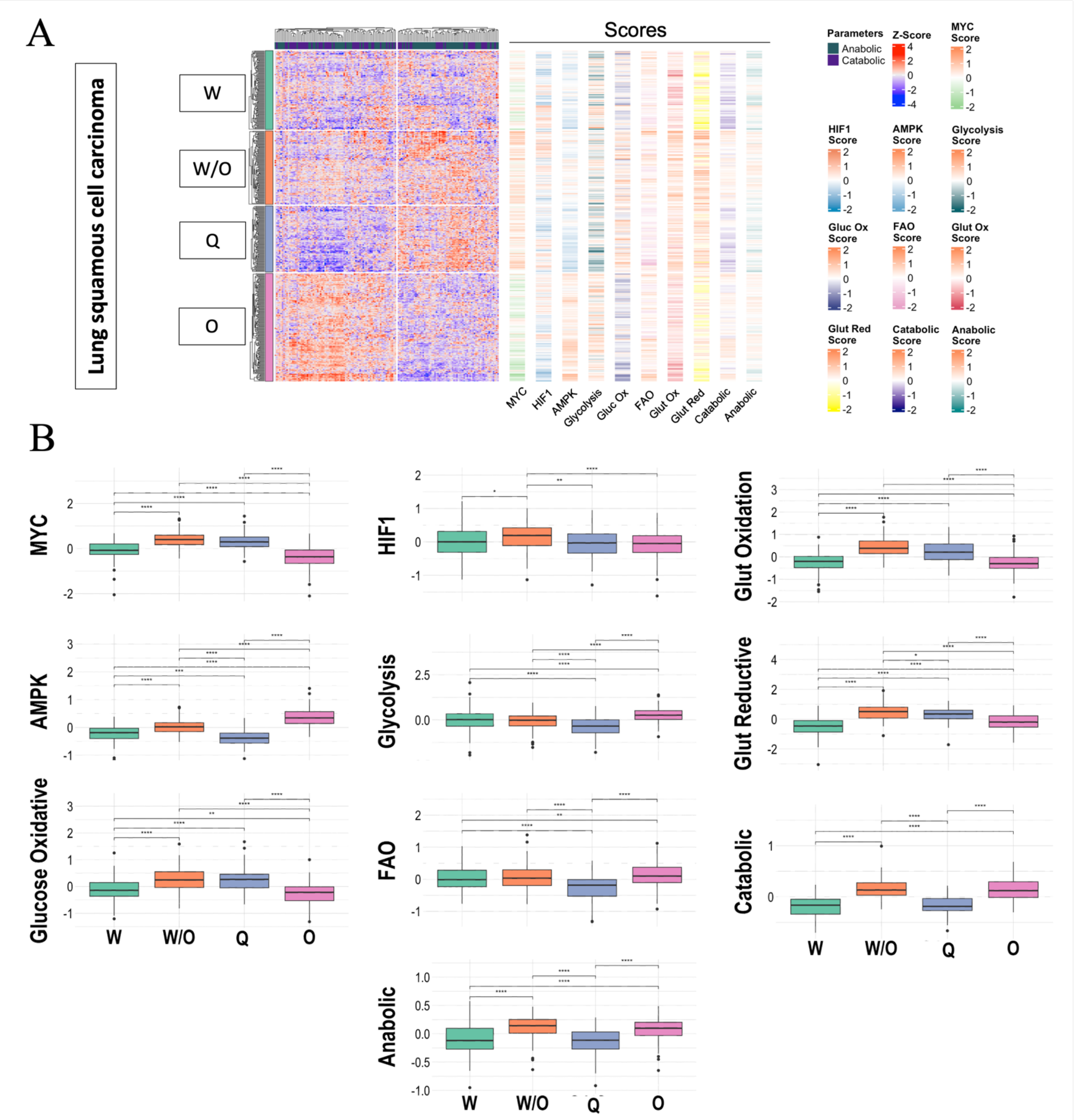


**Supplementary Figure 10. The association between gene activity and metabolic pathway activity for lung squamous cell carcinoma**. **A.** Heatmap of RNA-seq data. Each row represents a patient sample, and each column represents the expression of selected genes, which are divided according to their metabolic or anabolic activities. Four distinct clusters can be identified in the heatmap. The adjacent one-column heatmaps represent the scores for MYC, HIF-1, AMPK, Glycolysis, Glucose Oxidative, FAO, Glutamine Oxidative, Glutamine Reductive, Catabolic, and Anabolic activities, respectively. **B.** Box plots summarizing and showing the differences of the scores according to each identified cluster. A t-test was used to test the significance of each pair of clusters.

**Supplementary Figure 11**

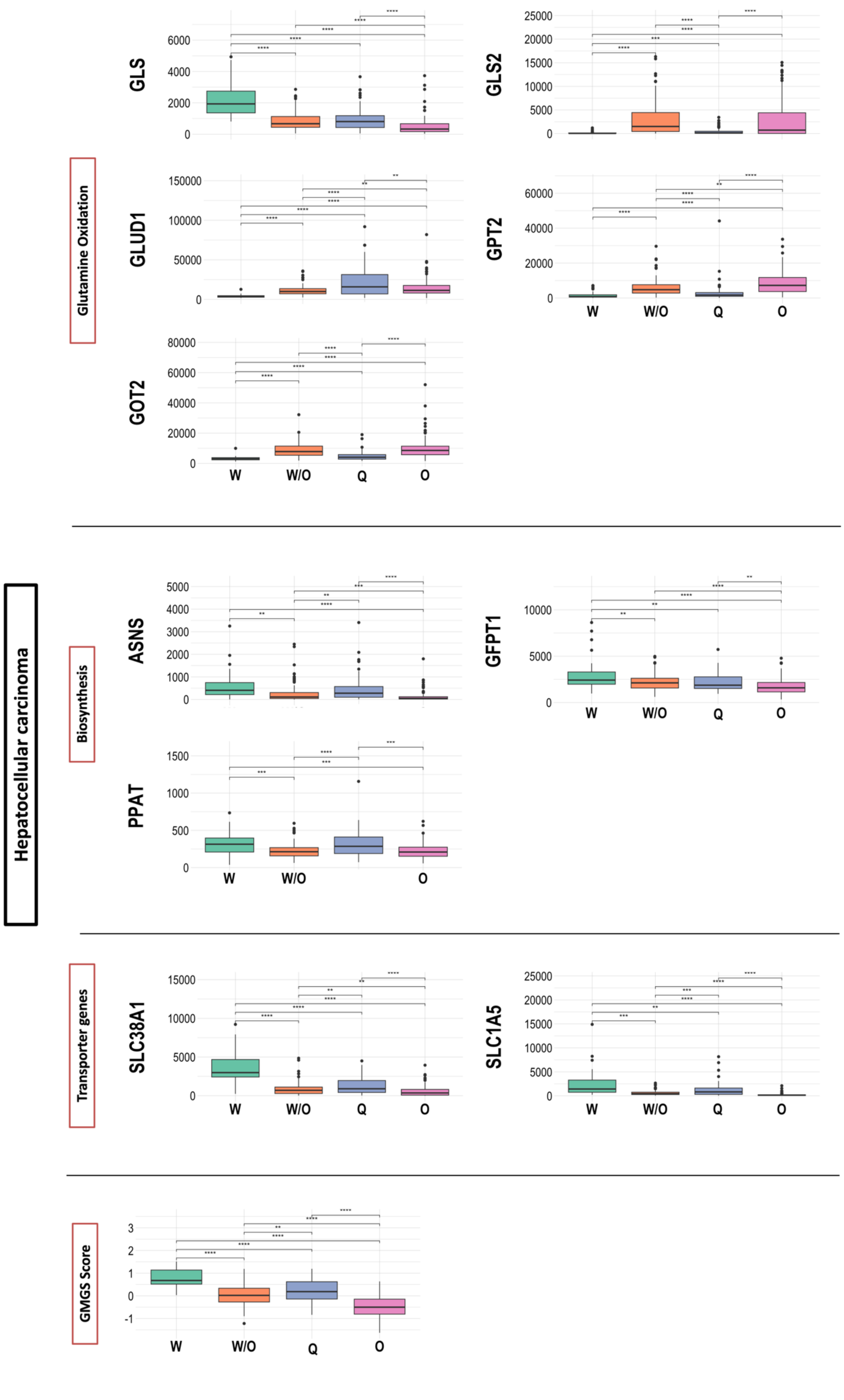

**Supplementary Figure 11. Comparison of glutamine metabolism relevant genes in the “O”, “W”, “W/O” and “Q” states in the hepatocellular carcinoma samples**. The glutamine metabolism relevant genes can be largely classified into three categories: oxidation genes (GLS, GLS2, GOT2, GLUD1 and GPT2), biosynthesis genes (ASNS, GFPT1, PPAT), and transporter genes (SLC38A1/2, SLC1A5), the gene signature (GMGS) (19) is also included. T-test was used to test the significance *, P < 0.05; **, P < 0.01; ***, P < 0.001.

### Supplementary Figure 12

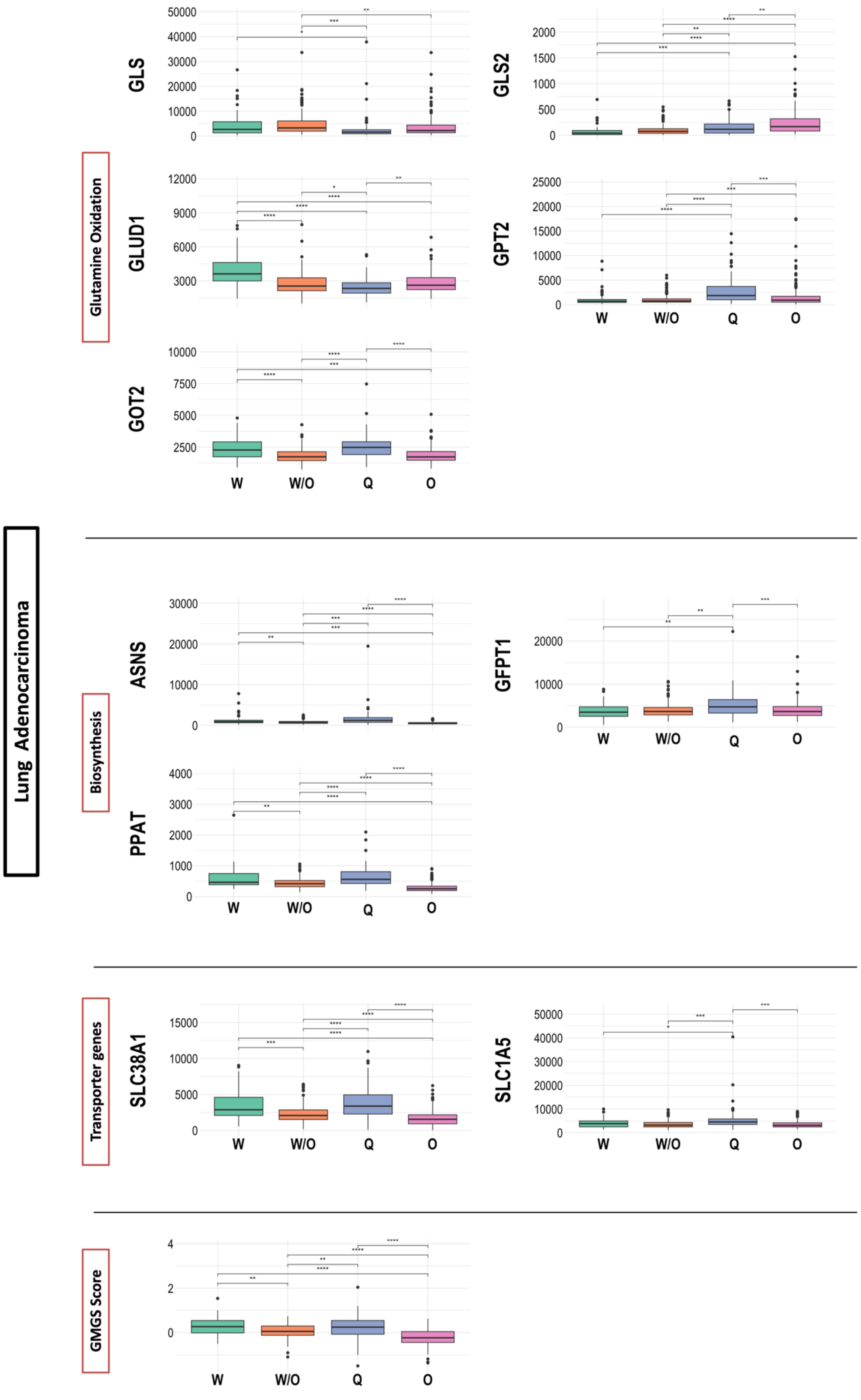

**Supplementary Figure 12. Comparison of glutamine metabolism relevant genes in the "O", "W", "W/O" and "Q" states in the Lung adenocarcinoma samples**. The glutamine metabolism relevant genes can be largely classified into three categories: oxidation genes (GLS, GLS2, GOT2, GLUD1 and GPT2), biosynthesis genes (ASNS, GFPT1, PPAT), and transporter genes (SLC38A1/2, SLC1A5), the gene signature (GMGS) (19) is also included. T-test was used to test the significance *, $P < 0.05$; **, $P < 0.01$; ***, $P < 0.001$.

**Supplementary Figure 13**

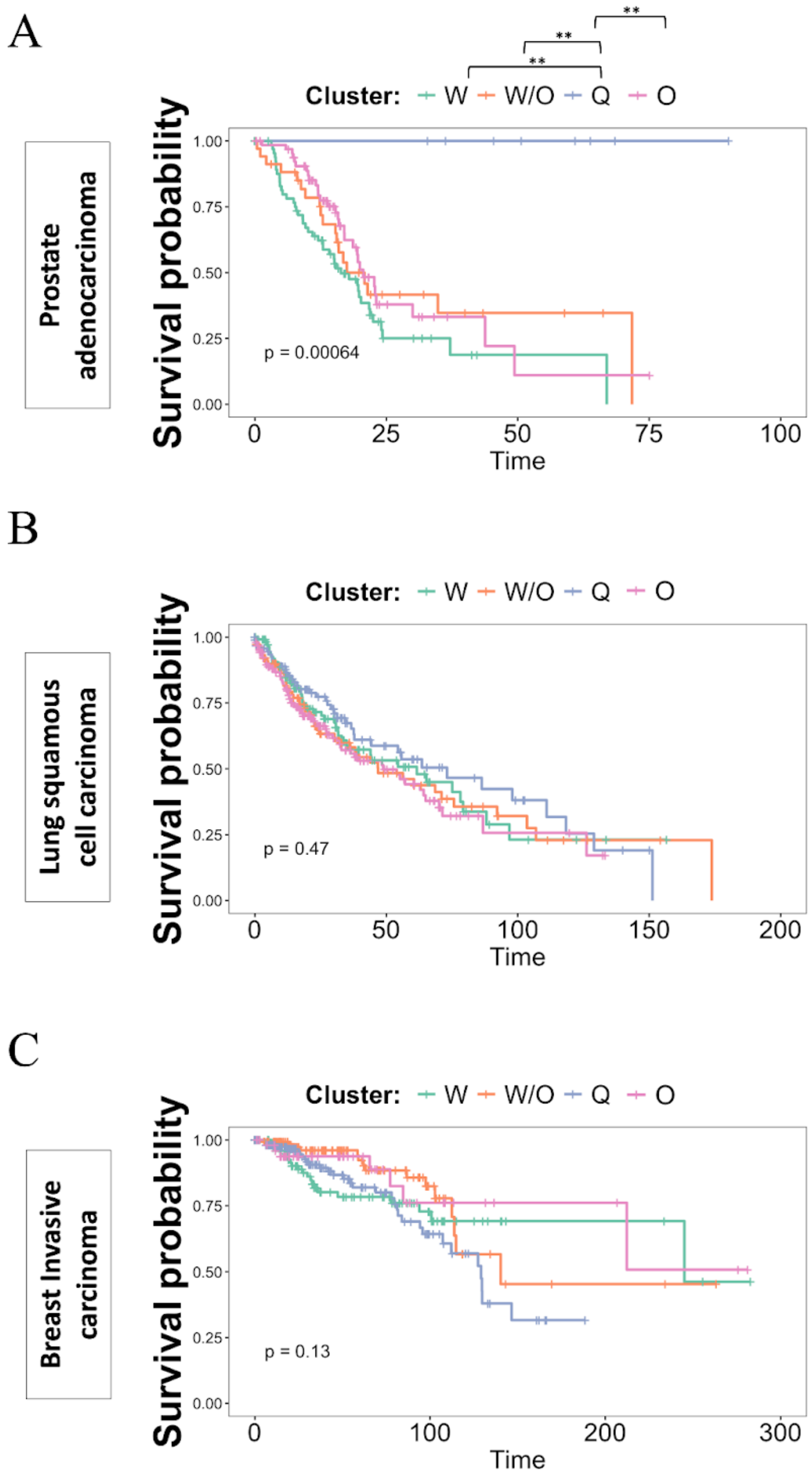


**Supplementary Figure 13 Survival curves stratified by cluster.** Survival curves for prostate adenocarcinoma (A), lung squamous cell carcinoma (B), and breast invasive carcinoma (C) datasets. Each dataset was separated in the four distinct phenotypes according to their metabolic or anabolic activities. The survival curves were estimated using the Kaplan-Meier method and compared using the log-rank test. Pairwise comparisons of survival distributions stratified by cluster are displayed above the survival curve for each pair of clusters. Significance levels are indicated as follows: *, $P < 0.05$; **, $P < 0.01$; ***, $P < 0.001$.

**Supplementary Table 1: Genes selected to evaluate metabolic pathway activity.**
List of genes selected in this study for various metabolic pathways. Gene sets were generated using Gene Ontology database AmiGO (AmiGO 2 version: 2.5.17, Carbon, S., et al 2009). Genes present in multiple pathways were excluded according to a priority list based on the order of appearance in the table.

| **Pathway** | **Genes used for the scoring metric** |
|---|---|
| AMPK (Yu, L. et al 2017) | ACADL, ACADM, ACOX1, ACOX3, ACSL1, ACSL3, ACSL4, ACSL5, ANGPTL4, APOC3, APOE, ATF4, ATF5, BAX, BCL2L11, BTG2, CAT, CCND1, CCND2, CLU, CPT1A, CPT1B, CPT2, CREM, CYP27A1, CYP4A11, CYP7A1, DDB1, DGAT1, DNMT1, EGR1, EHHADH, ELN, FADS2, FASLG, FOS, FOSB, FOSL1, FOXA2, G6PC, G6PC2, G6PC3, GADD45A, GADD45B, GADD45G, GK, HES1, HNF4A, HSPD1, ID1, ID3, JUNB, JUND, KLF5, LPL, MAFA, MAPK9, MECR, MMP9, NCOA2, NR1H3, NR4A1, NR4A2, NUP155, NUP88, NUP98, OLR1, ONECUT1, ONECUT2, PCK1, PCK2, PDK4, PDX1, PRMT1, PRMT3, RUVBL1, SCD, SLC2A4, SOD2, SORBS1, SP1, STAT5A, TOB1, TXNIP. |
| HIF-1 (Yu, L. et al 2017) | ADM, ALDH4A1, ALDOA, ALDOC, ARHGEF1, ASPH, BHLHE40, BNIP3, BTAF1, CA9, CCNB1, CITED2, CP, DDIT4, EDN1, EGLN1, EGLN3, ENO1, EPRS, ETS1, FN1, GLCCI1, GRK6, HACD3, HSP90B1, IVNS1ABP, KDM3A, MECOM, MET, MXD1, MYLK, NAMPT, NDRG1, NOS2, NOS3, P4HA1, P4HA2, P4HTM, PDGFA, PFKL, PGAM1, PGK1, PLOD1, RARA, RBPJ, RNF165, RRAGD, RSBN1, SERPINE1, SLC31A1, SLC7A6, SOX6, SSRP1, STC2, TFRC, TGFB3, TMEFF1, TMEM45A, VEGFA. |
| Glucose Oxidation (G1) (Jia, D. et al 2019) | PDHA1, ACO2, IDH1, OGDH, SDHA, SDHC, FH, MDH1, CS, PC |
| Glycolysis (G2) (Jia, D. et al 2019) | HK1, GPI, PFKM, ALDOA, TPI1, GAPDH, PGK1, PGAM2, ENO1, PKM |
| FAO (F1) (Jia, D. et al 2019) | CD36, ACSL1, SLC25A20, CPT1, CPT2, ACADVL, ACADL, ACADM, ACADS, ACADSB, ACAD8, ACAD9, ACAD10, ECHS1, HADH, ACAA1, ACAA2 |
| MYC | MTDH, ACTL6A, ENO1, BAX, BCAT1, COMMD3-BMI1, POLR3D, CAD, CDC25A, CDK4, CCNB1, CCND2, DDX18, DKC1, E2F3, EIF2S1, EIF4A1, EIF4E, EIF4G1, FOSL1, GAPDH, KAT2A, SLC2A1, GPAM, HUWE1, HMGA1, HSPD1, HSPA4, HSP90AA1, ID2, IREB2, CDCA7, KIR3DL1, LDHA, LIN28B, MAX, NDUFAF2, RIOX2, MMP9, MTA1, MYC, MYCT1, NBN, NME1, NME1-NME2, PMAIP1, NPM1, NCL, ODC1, TP53, PDCD10, PEG10, PFKM, PIM1, PRDX3, PTMA, RCC1, RPL11, SERPINI1, SHMT1, SMAD3, SMAD4, SNAI1, SUPT3H, SUPT7L, BIRC5, TAF10, TAF12, TAF4B, TAF9, TERT, TFRC, RUVBL1, RUVBL2, KAT5, TK1, TRRAP, UBTF |

| | |
|---|---|
| Reductie glucose (Gre) | G6PC2, G6PC3, G6PC1, PCK2, PCK1, PC, GPI, TPI1, FBP1, FBP2 |
| Glutamine oxidation (Q1) | GLS, GLS2, GOT1, GOT2, GLUD1, GPT2 |
| GSH synthesis (QSH) | GGT1, GGT7, GGT5, GSS, OPLAH, CNDP2, GGCT, GGT6 |
| Reductive glutamine (Qre) | GLUD2, QARS1, GCLC, CPS1, GCLM, EPRS1, CAD, GLUL, ABAT, GAD2, GAD1, PPAT, ASNS, GFPT1 |
| Reductive Fatty acids (Fre) | PCCB, DEGS1, MECR, SCD, PLA2G1B, ABHD3, ACACB, ACSF3, PRKAA2, PTGS2, PTGES, FA2H, FAAH, GSTM4, CYP1A1, MGLL, HTD2, HSD17B8, AKR1C3, FASN, LIPE, ABCD1, PRKAG2, CBR4, ELOVL6, ACACA, MCAT, DECR2, PECR, HPGDS, FADS1, PRKAB2, CBR1, PRXL2B, ACLY, NDUFAB1, PLA2G10, OLAH, ALOX12B, FADS2, ALOX5, LTC4S, ALOX12, ELOVL2, ALOX15B, ELOVL7, PTGDS, PLA2G4A, ALOXE3, ALOX15, TBXAS1, SCD5, PTGS1 |
| NADPH oxidase-derived ROS (Rn) | CYBB, CYBA, NCF1, NCF2, NCF4, NOX1, NOX3, NOX4, NOX5 |
| Mitochondrial ROS (Rm) | NDUFS1, CYTB, SDHA, MAOB, CYCS, VDAC, UCP2, SOD2, CAT, GPX1 |
| Glutamine Uptake (Q0) | SLC25A22, SLC38A5, SLC6A19, SLC7A5, SLC38A2, SLC38A1, SLC25A13, SLC1A5, SLC38A3, SLC38A8, SLC7A7, SLC7A6, SLC6A14, SLC7A11, SLC38A7 |
